# DNA damage independent inhibition of NF-κB transcription by anthracyclines

**DOI:** 10.1101/2020.04.27.065003

**Authors:** Ângelo Ferreira Chora, Dora Pedroso, Eleni Kyriakou, Nadja Pejanovic, Henrique Colaço, Raffaella Gozzelino, André Barros, Katharina Willmann, Tiago Velho, Catarina F. Moita, Isa Santos, Pedro Pereira, Sílvia Carvalho, Filipa Batalha Martins, João A. Ferreira, Sérgio Fernandes de Almeida, Vladimir Benes, Josef Anrather, Sebastian Weis, Miguel P. Soares, Arie Geerlof, Jacques Neefjes, Michael Sattler, Ana C. Messias, Ana Neves-Costa, Luís Ferreira Moita

**Affiliations:** Instituto de Medicina Molecular, Faculdade de Medicina, Universidade de Lisboa, 1649-028 Lisboa, Portugal; Innate Immunity and Inflammation Laboratory, Instituto Gulbenkian de Ciência, Rua da Quinta Grande 6, 2780-156 Oeiras, Portugal; Institute of Structural Biology, Helmholtz Zentrum München, Ingolstädter Landstr. 1, 85764 Neuherberg, Germany; Biomolecular NMR and Center for Integrated Protein Science Munich at Department Chemistry, Technical University Munich, Lichtenbergstr. 4, 85747 Garching, Germany; Chronic Diseases Research Center (CEDOC), NOVA Medical School (NMS), Lisbon, Portugal; Centro Hospitalar Lisboa Norte - Hospital de Santa Maria, EPE, Avenida Professor Egas Moniz, 1649-035, Lisbon, Portugal; Serviço de Cirurgia, Centro Hospitalar de Setúbal, Setúbal, Portugal; EMBL Genomics Core Facilities, D-69117 Heidelberg, Germany; Feil Family Brain and Mind Research Institute, Weill Cornell Medicine, 407 East 61st Street RR409, New York, NY, 10065, USA; Institute for Infectious Disease and Infection Control, Jena University Hospital, 07747 Jena, Germany; Department of Anesthesiology and Intensive Care Medicine, Jena University Hospital, 07747 Jena, Germany; Center for Sepsis Control and Care, Jena University Hospital, 07747 Jena, Germany; Inflammation Laboratory, Instituto Gulbenkian de Ciência, Rua da Quinta Grande 6, 2780-156 Oeiras, Portugal; Department of Cell and Chemical Biology, LUMC, Leiden, The Netherlands; Instituto de Histologia e Biologia do Desenvolvimento, Faculdade de Medicina da Universidade de Lisboa, 1649-028 Lisboa

**Author notes:** These authors contributed equally (Ângelo Ferreira Chora, Dora Pedroso). Correspondence: Ana Neves-Costa, and Luis F. Moita.

**Keywords:** Anthracyclines, nuclear factor kappa B (NF-κB), inflammation, DNA damage

## Abstract

Anthracyclines are among the most used and effective anticancer drugs. Their activity has been attributed to DNA double-strand breaks resulting from topoisomerase II poisoning and to eviction of histones from select sites in the genome. Here we show that the extensively used anthracyclines Doxorubicin, Daunorubicin and Epirubicin, decrease the transcription of nuclear factor kappa B (NF-κB)-dependent gene targets, but not interferon responsive genes. Using an NMR-based structural approach, we demonstrate that anthracyclines disturb the complexes formed between the NF-κB subunit RelA and its DNA binding sites. The variant anthracyclines Aclarubicin, Doxorubicinone and the newly developed Dimethyl-doxorubicin, which share anticancer properties with the other anthracyclines but do not induce DNA damage, also suppressed inflammation, thus uncoupling DNA damage from the effects on inflammation. These findings have implications for anticancer therapy and for the development of novel anti-inflammatory drugs with limited side effects for life-threatening conditions such as sepsis.

## Introduction

Inflammation is critical to homeostasis maintenance. Recognition of microbial-associated molecular patterns (MAMPs) and non-microbial molecules by specialized sensors is a key first step for inflammatory responses. Engagement of germline-encoded pattern recognition receptors (PRRs), triggers signal transduction pathways leading to the activation of transcription factors controlling the expression of a large number of pro-inflammatory genes (Medzhitov and Horng, 2009).

The nuclear factor kappa B (NF-κB) family of transcription factors, comprising p65/RelA, RelB, c-Rel, p100 and p105, plays a prominent role in regulating inflammation (Hayden and Ghosh, 2008). All members share the N-terminal REL-homology domain (RHD) for IκB (inhibitor of κB) interaction, nuclear localization, dimerization and DNA binding. The C-terminal transactivation domain (TAD) is required for transcriptional activity (Chen and Greene, 2004). While NF-κB activation is strictly required to trigger the inflammatory responses while preventing cytotoxicity, sustained NF-κB activity can perpetuate inflammation, contributing to the pathogenesis and establishment of non-transmissible, often chronic pathological conditions (Baldwin, 2001).

At steady state, transcriptionally active NF-κB family members are retained in the cytoplasm through interactions with IκB molecules. RelA is the most abundant representative member of this family. IκBα phosphorylation in response to a variety of stimuli leads to its proteasomal degradation and to RelA nuclear translocation as a consequence, ultimately culminating in DNA binding and transcription of NF-κB regulated genes (Chen and Greene, 2004, Ghosh and Baltimore, 1990). RelA-dependent IκBα re-synthesis is essential for RelA nuclear eviction and the timely termination of NF-κB transcriptional activity (Sun et al., 1993, Beg and Baldwin, 1993). Posttranslational modifications within both RHD and TAD domains add an additional layer of regulation over NF-κB-dependent gene transcription by controlling the association with IκB, nuclear localization, interaction with transcriptional cofactors, DNA binding and stability (Chen and Greene, 2004, Huang et al., 2010).

Anthracyclines have been used as anticancer drugs for many decades and Doxorubicin (Doxo), in particular, is used in therapies against a wide variety of cancers (reviewed in (Hande, 1998)). The cytotoxicity of anthracyclines has been mostly attributed to their ability to target and inhibit topoisomerase II (TopoII) (Tewey et al., 1984). In rapidly replicating cells, anthracyclines stabilize the TopoII-DNA cleavable complex, potentiating DNA double-strand breaks, activation of DNA damage responses (DDRs), and ultimately senescence or programmed cell death by apoptosis (Nitiss, 2009) (Eom et al., 2005). Other clinically relevant biological activities of anthracyclines include DNA intercalation, helicase inhibition and free radical formation, but their mechanisms of action are still incompletely understood (Hande, 1998). More recently, anthracyclines were shown to induce histone eviction from discrete chromosomal regions and to contribute to apoptosis in a TopoII-independent manner (Pang et al., 2013) (Yang et al., 2013). Cytotoxicity due to histone eviction was observed in AML blasts, highlighting the potential clinical impact of these other less exploited activities of anthracyclines (Pang et al., 2013).

We have previously shown that anthracyclines strongly suppress cytokine secretion in THP-1 cells challenged with pro-inflammatory stimuli and in mouse models of sepsis (Figueiredo et al., 2013), a finding recently corroborated in primary human macrophages (Köse-Vogel et al., 2020). In such models, low-dose anthracyclines induce disease tolerance leading to less severe sepsis with decreased mortality independently of the pathogen load (Figueiredo et al., 2013). The DDR and the DNA damage sensor ATM (Ataxia Telangiectasia Mutated) in particular are required for anthracycline-mediated induction of disease tolerance to bacterial infections (Figueiredo et al., 2013), and Epirubicin (Epi) was shown to induce and activate ATM in breast cancer cells (Millour et al., 2011). However, the mechanism whereby anthracyclines regulate cytokine production remains obscure (Neves-Costa and Moita, 2017). Although the role of ATM in DDR has been extensively documented (comprehensively reviewed in (Maréchal and Zou, 2013)), ATM participates in a complex network of signaling pathways including activation of NF-κB (Huang et al., 2010, Piret et al., 1999, Li et al., 2001, Wu et al., 2006).

Here we demonstrate that anthracyclines modulate the inflammatory response in primary macrophages by interfering directly with RelA DNA binding independently of the DDR and in a manner that contributes to the resolution of inflammation. The use of various anthracycline variants allowed the uncoupling of this effect from the classical DNA damage response and hints to a direct interaction and inhibition of RelA binding to DNA.

## Results

### Cytokine secretion and DNA damage

To investigate the role of DNA damage responses (DDRs) in cytokine downregulation induced by anthracyclines (Figueiredo et al., 2013), we measured cytokine concentrations in conditioned media in response to Epirubicin (Epi) upon activation of WT and *Atm*-deficient bone marrow-derived mouse macrophages (BMDMs) by pro-inflammatory stimuli. Pre-treatment of BMDMs with Epi downregulated cytokines following *E. coli* challenge independently of ATM, with Epi leading to a dose-dependent decrease in TNF, IL12, IL6 and Cxcl10 production not only in WT, but also in *Atm*^-/-^ macrophages (Fig. 1A). Upon activation, WT and *Atm*^-/-^ BMDMs secreted comparable amounts of cytokines (Supplementary Fig. 1A). We then tested cytokine secretion in the presence of the ATM inhibitor Ku-55933 and confirmed that ATM is dispensable for limiting TNF and IL12 secretion by Epi (Fig. 1B). At the doses used Epi was not cytotoxic (Supplementary Fig. 1B) nor did it activate cytokine secretion per se.

**Figure 1.**
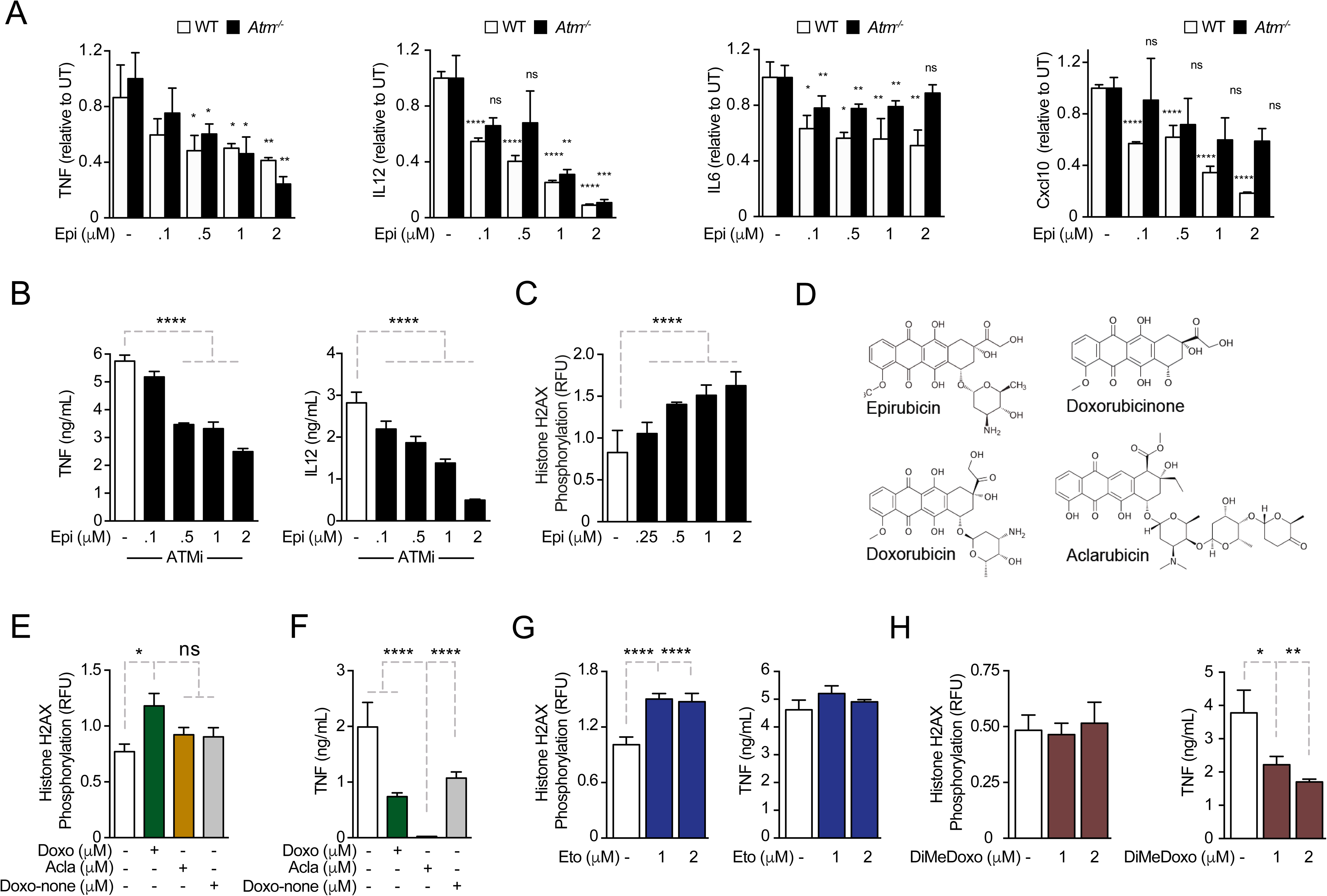
Anthracyclines regulate cytokine secretion independently of ATM. **A**, Cytokine secretion of TNF, IL12, IL6 and Cxcl10 was quantified by ELISA following *E. coli* challenge in the presence of various doses of Epirubicin (Epi) in WT and *Atm*-/- macrophages; **B**, Secretion of TNF and IL12 was quantified in macrophages treated with various doses of Epi and the ATM inhibitor KU-55933; **C**, H2AX phosphorylation was quantified by ELISA in Epi-treated macrophages, normalized to total H2AX and shown as relative fluorescence intensities (RFU); **D**, Schematic representation of the molecular structures of the anthracyclines Epi, Aclarubicin (Acla), Doxorubicin (Doxo) and Doxorubicinone (Doxo-none); **E**, H2AX phosphorylation was quantified in the presence of Doxo, Acla and Doxo-none; **F**, TNF secretion was quantified in the presence of Doxo, Acla and Doxo-none; **G**, **H**, H2AX phosphorylation and TNF secretion were quantified in macrophages treated with Etoposide (Eto) and Dimethyl-doxorubicin (diMe-Doxo) following *E. coli* challenge. The assays show arithmetic means and standard deviations of technical replicates from one representative animal of at least three independent animals tested. p < 0.05 (*); p < 0.01 (**); p < 0.001 (***); p < 0.0001 (****).

As cytokine downregulation by Epi was not ATM dependent, we hypothesized that the DDR was not required for Epi-dependent cytokine suppression. We started by confirming, using the alkaline comet assay to detect strand breaks, that DNA damage caused by Epi is time- and dose-dependent (Supplementary Fig. 1C) and then we proceeded with DNA damage quantification using phosphorylation of histone H2AX at Ser139 (γH2AX) as a surrogate marker. In BMDMs, DNA damage was strongly induced by Epi in the range of concentrations that modulated cytokine production (Fig. 1C) and was comparable to that of Etoposide (Eto), another well-studied TopoII inhibitor that causes DNA breaks and induces ATM-mediated DDRs (Supplementary Fig. 1D) (Caporossi et al., 1993, Banáth and Olive, 2003). γH2AX levels also showed that the inflammatory challenge did not induce significant extra DNA damage in the conditions used (Supplementary Fig. 1E). The lack of interdependency between DNA damage and cytokine modulation was then assayed for other anthracyclines. These drugs all share a tetracycline ring decorated with one or multiple amino sugars (Fig. 1D) and their activities range from failing to induce DNA breaks to being potent inducers of double strand breaks (Pang et al., 2013). Eto is not a member of the anthracycline class but inhibits TopoII like Doxo and Epi. DNA damage caused by anthracyclines was also diverse in BMDMs: whereas Epi and the closely related Doxorubicin (Doxo) led to a dose-dependent increase in γH2AX signal, Aclarubicin (Acla) and Doxorubicinone (Doxo-none) did not induce significant damage (Fig. 1C and 1E). The comet assay confirmed that DNA damage caused by Acla is comparable to the basal damage levels in untreated cells, in sharp contrast to the highly damaging effects of Epi and Eto (Supplementary Fig. 1F). Acla and Doxo-none were then tested for their ability to regulate cytokines. Acla led not only to the most pronounced downregulation of TNF, but also of all the other cytokines tested at concentrations without significant toxicity (Supplementary Fig. 2A,B). Whereas downregulation of TNF and IL12 was a common property of the anthracyclines tested, Doxo-none was less potent than Epi, Doxo and Acla (Fig. 1F and compare with 1A). Doxo-none does not induce DNA breaks nor histone eviction (Qiao et al., 2020). To further uncouple the DDR from cytokine modulation by anthracyclines, we tested if Eto and a variant of Doxo that was made incapable of causing DNA damage while leaving histone eviction activity - but retains a cytotoxic profile similar to Doxo by the introduction of 2 methyl groups in the sugar ring, Dimethyl-doxorubicin (diMe-Doxo) (Qiao et al., 2020) - could also downregulate cytokines. Figures 1G and 1H show that TNF was not downregulated by Eto but was by diMe-Doxo. In line with this observation, none of the other anthracyclines tested regulated cytokine production differently in *Atm*^-/-^ BMDMs or in the presence of the ATM inhibitor (Supplementary Fig. 2C, D). Therefore, structural damage to DNA alone is unlikely to explain the observed effect of anthracyclines on cytokine secretion. The histone eviction activity of anthracycline drugs is more associated to the control of cytokine secretion upon LPS challenge.

### Anthracyclines negatively regulate the transcription of NF-**κ**B target genes

Before dissecting the molecular mechanisms of cytokine downregulation by anthracyclines, we investigated the *in vivo* effects of the administration of these drugs in a model of septic shock. To this end, we co-administered Epi or Acla to mice that were challenged with LPS. We found that the circulating levels of TNF were significantly decreased in mice that we treated with either Epi or Acla, 6 h following the initial challenge (Fig. 1A), suggesting that our findings have *in vivo* physiological relevance. We then investigated the effects of anthracyclines on transcription and their dependence on DNA damage, we performed RNA sequencing (RNAseq) in BMDMs pre-treated with either Epi or Acla and challenged thereafter with LPS (Supplementary Fig. 3A). By comparing mRNA levels of untreated with that of 4h LPS-treated BMDMs, we detected strong induction of pro-inflammatory gene expression, in line with the well-described patterns of transcriptional regulation in response to this TLR4 agonist (Fig. 2B, lane 1). Differential expression analysis revealed 455 genes at least 5-fold upregulated in LPS-treated BMDMs compared with untreated, whereas only 75 genes were downregulated to the same extent by LPS (Supplementary Fig. 3B). The transcriptomes also pointed to specific gene expression signatures by Epi and Acla, partly overlapping but not identical, both in unstimulated BMDMs (Fig. 2B, lanes 2 and 3) and in the presence of LPS (Fig. 2B, lanes 4 and 5). This could be the result of different genomic location preferences between these drugs, as shown for Acla vs Daunorubicin (Pang et al., 2015). As anticipated from the cytokine secretion results, repression of target genes by Acla was stronger than by Epi (Fig. 2B, compare lanes 4 and 5). Functional enrichment analysis showed that both Epi and Acla repress cytokine production, amongst other effector functions of macrophages (Supplementary Table 1). The fact that both Epi and Acla regulate inflammatory gene expression also supports a DNA damage-independent mechanism. By quantitative RT-PCR, we confirmed downregulation of TNF, IL12, IL6 and Cxcl10 by both Epi and Acla (Supplementary Fig. 3C), and also of a broader subset of pro-inflammatory mediators, including IκBα (Fig. 2C). We then searched for promoter motifs in repressed genes (Chang and Nevins, 2006) and, not surprisingly, we detected an NF-κB signature shared by genes mutually downregulated by Epi and Acla. To compare with NF-κB-dependent genes, we extended the RT-PCR analysis to other TLR4-induced transcriptional programs, namely IRF-dependent interferon transcription (Kawai and Akira, 2007). IRF3-dependent IFNβ (*Ifnb1*) transcription was strongly induced by *E. coli* as expected, but not downregulated by Epi or Acla (Fig. 2D). In agreement, IFNβ secretion was also not suppressed by the anthracyclines tested (Supplementary Fig. 3D). IFNα (*Ifna1, Ifna4*) transcription, typically mediated by virally induced receptors such as TLRs7/9, was not induced, nor was the expression of these genes further downregulated by Epi or Acla (Supplementary Fig. 3E). In combination, these results indicate that anthracyclines do not compromise the overall cellular transcription. Instead, the transcriptional profiles point to a negative effect of Epi and Acla specifically on NF-κB-regulated gene expression. This is compatible with previous observations that the anthracyclines Doxo and Daunorubicin (Dauno) repress TNF-induced NF-κB transactivation in cancer cells (Campbell et al., 2004) (Ho et al., 2005).

**Figure 2.**
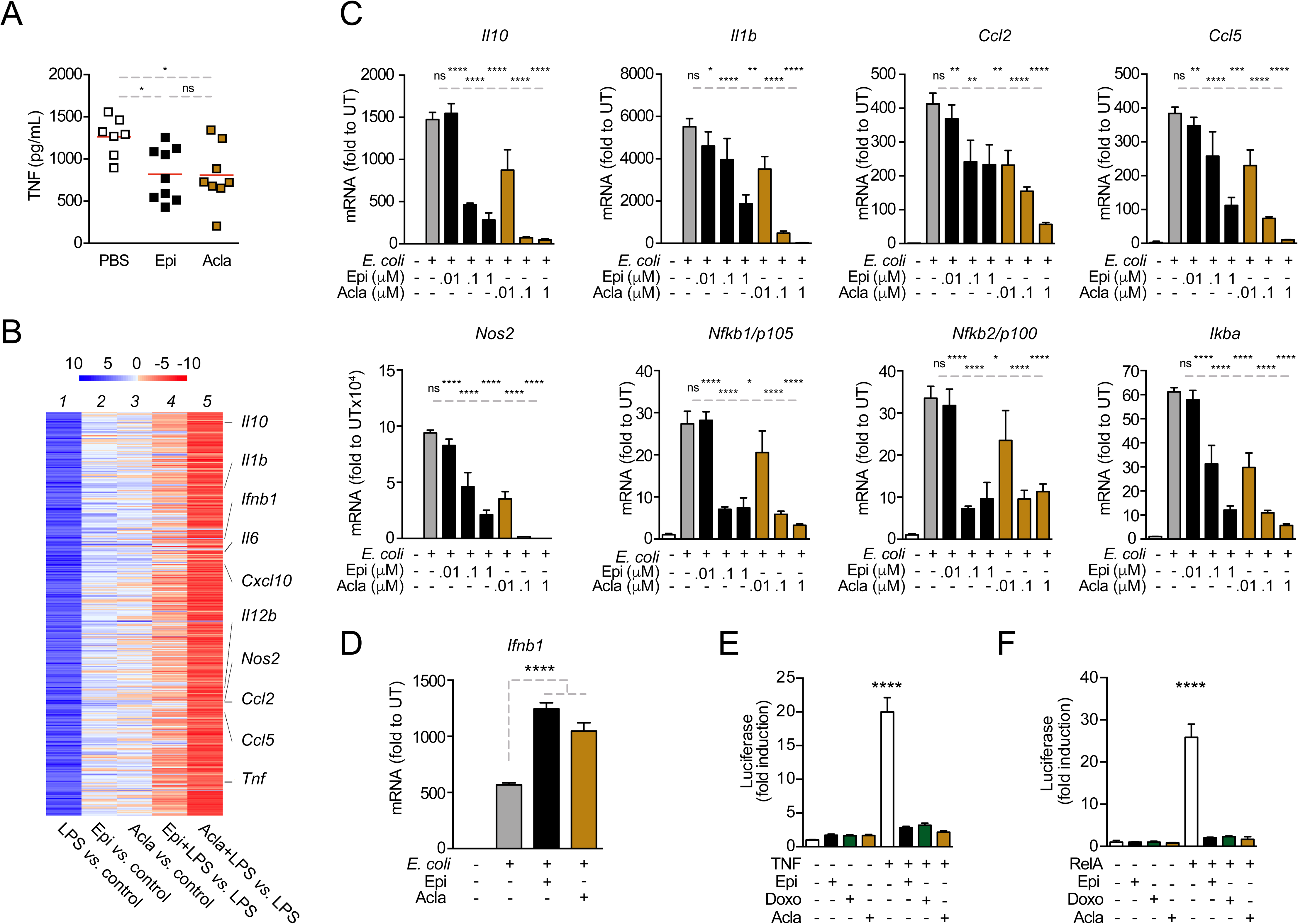
Anthracyclines regulate NF-*κ*B-dependent transcription. **A**, TNF concentrations in serum 8h post *E. coli* challenge in mice treated with PBS, Epirubicin (Epi) or Aclarubicin (Acla). **B,** RNAseq in macrophages stimulated with LPS for 4h and treated with Epi or Acla; **C**, Gene expression was quantified by quantitative RT-PCR in macrophages following *E. coli* challenge and treated with various doses of Epi and Acla; results were normalized to expression in untreated conditions (UT); **D**, Gene expression of *Ifnα* was quantified as in B; **E**, Luciferase quantification of a κB reporter in HEK293 cells stimulated with TNF and treated with 50nM of Epi, Doxorubicin (Doxo) and Acla; **F**, Luciferase quantification of a *κ*B reporter in HEK293 cells treated with 2µM of Epi, Doxo and Acla in the absence or presence of a vector overexpressing RelA. **B**, **C** and **D** show arithmetic means and standard deviations of technical replicates from one representative animal of at least three independent animals tested. p < 0.05 (*); p < 0.01 (**); p < 0.001 (***); p < 0.0001 (****).

To test whether anthracyclines affect mRNA stability of pro-inflammatory genes, we treated BMDMs with actinomycin D (ActD) 2h after stimulation in the presence or absence of Epi and measured the rate of mRNA decay. We did not observe significant differences in the half lives of TNF mRNAs between non-treated and Epi-exposed BMDMs (Supplementary Fig. 3F). The effect of anthracyclines on NF-κB-dependent gene expression is therefore likely to be caused by changes in transcription and not due to regulation of mRNA stability.

To further address transcriptional regulation by anthracyclines, we used a NF-κB luciferase reporter assay. In transiently transfected HEK293 cells stimulated with TNF, Epi, Doxo, Acla and Dauno effectively repressed reporter expression to basal unstimulated levels (Fig. 2E). To search for additional evidence for a role of anthracyclines in suppressing NF-κB-dependent transcription, we co-transfected the NF-κB luciferase reporter together with a vector expressing the full-length NF-κB family member RelA. Whereas RelA considerably stimulated luciferase activity as expected, anthracyclines markedly inhibited the expression of the reporter (Fig. 2F). This suggests that anthracyclines act downstream of IκBα, targeting RelA directly or indirectly.

### Epirubicin affects NF-**κ**B sub-cellular localization

NF-κB nuclear translocation is required for DNA binding and initiation of transcription (Ghosh and Baltimore, 1990). We tested whether the downregulation of NF-κB targets by anthracyclines was due to impaired RelA nuclear translocation. RelA nuclear levels in untreated BMDMs rapidly increased following inflammatory challenge and RelA slowly relocated overtime, being mostly cytoplasmic 4h after stimulation (Fig. 3A, control panel). In BMDMs exposed to Epi, RelA translocated to the nucleus upon *E. coli* challenge, but despite decreased NF-κB-dependent gene expression, RelA remained nuclear at all time points analyzed (Fig. 3A, Epi panel). Proteolytic degradation and re-synthesis of the NF-κB inhibitor IκBα is a central mechanism controlling the sub-cellular localization of NF-κB factors (Ghosh and Baltimore, 1990; Sun et al., 1993; Beg and Baldwin, 1993). From our mRNA analysis, we expected Epi to affect IκBα cellular levels (Fig. 2B), and therefore we decided to test the effects of Epi on IκBα protein throughout time. As extensively reported, IκBα was degraded and protein levels restored to initial values within 60 minutes in non-treated BMDM. In contrast, Epi pre-treatment profoundly diminished IκBα cellular levels throughout the time course (Fig. 3B), as anticipated form our previous results. This regulation was not stimulus-dependent, as was also observed upon TNF stimulation (Supplementary Fig. 3G). Reduced stimulus-induced IκBα synthesis was reported to promote nuclear localization of RelA, an effect not associated with increased NF-κB transcriptional activity (Hochrainer et al., 2007).

**Figure 3.**
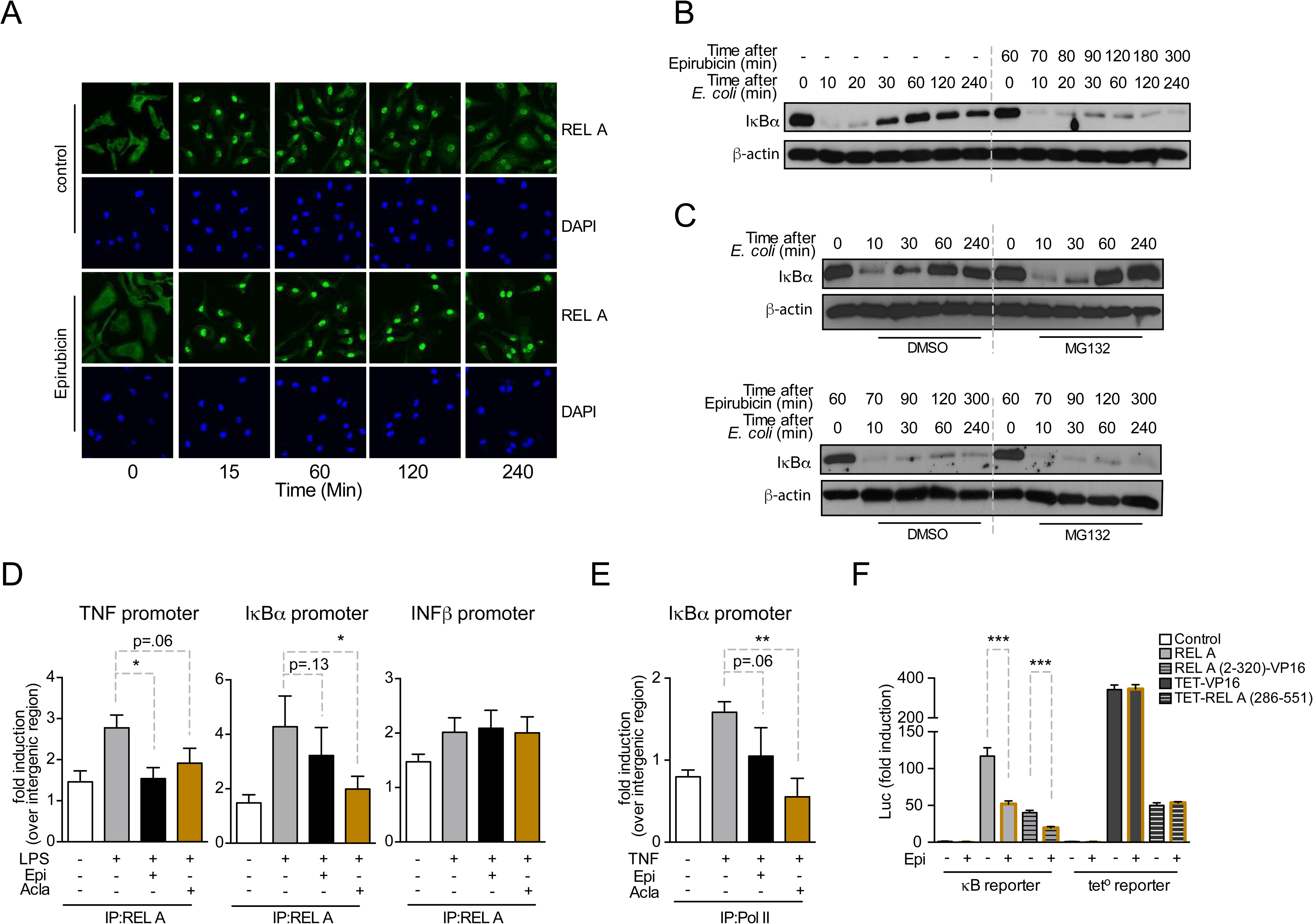
Anthracyclines affect RelA sub-cellular localization. **A**, RelA immunolocalization in macrophages challenged with *E. coli* and left untreated (control) or treated with 2µM of Epirubicin (Epi); **B**, I*κ*B*α* degradation kinetics in macrophages following *E. coli* challenge in the absence or presence of 2µM of Epi for 1h at the time of *E. coli* challenge; **C**, Macrophages were either left untreated (top panel) or treated with Epi (bottom panel) and challenged with *E. coli* for the indicated times in the presence of the proteasome inhibitor MG132 or its vehicle DMSO; **D**, Macrophages were challenged with LPS, treated with 2µM of Epi or Aclarubicin (Acla) and an antibody anti-RelA was used to immunoprecipitate the associated chromatin, from where the promoter sequences of *IκBα, Tnf* and *Ifnβ* were amplified; **E**, HEK293 cells were challenged with TNF, treated with 2µM of Epi or Acla and an antibody anti-PolII was used to immunoprecipitate the associated chromatin, from where the promoter sequence of *IκBα* was amplified; **F**, HEK293 cells were transiently transfected with the κB-luc reporter alone or in conjunction with full length RelA or RelA (2-320)-VP16 or TET-RelA (268-551) and cells were left untreated or treated with 2µM of Epi for 16h.

A role for Doxo in proteasome activation has been proposed (Liu et al., 2008). To test if increased proteasome-dependent degradation also contributes to low IκBα levels, we examined IκBα degradation kinetics upon inhibition of the proteasome. While blockage of proteasome activity by MG-132 increased IκBα protein levels upon *de novo* synthesis following *E. coli* challenge (Fig 3C, upper panel), IκBα in cells pre-exposed to Epi never recovered (Fig 3C, lower panel). It is therefore improbable that IκBα regulation by Epi is due to any role for the anthracyclines in proteasomal activity. Instead, the role of Epi in repressing *E. coli*-driven IκBα mRNA expression (Fig. 2B) is likely the reason for RelA nuclear localization observed upon anthracycline pre-treatment. Our data suggests that anthracyclines are capable of breaking the critical negative feedback loop that maintains the cellular responsiveness to subsequent inflammatory stimuli.

### Epirubicin and Aclarubicin regulate NF-**κ**B binding to its targets

We further investigated by chromatin immunoprecipitation (ChIP) how anthracyclines suppress NF-κB-dependent pro-inflammatory programs. Firstly, we confirmed that the effects of anthracyclines in BMDM could be replicated in HEK293 cells (Supplementary Fig. 4). ChIP at the promoter of selected NF-κB target genes in HEK293 cells revealed that RelA binding following inflammatory activation was weakened by pre-treatment with Epi and Acla, ranging from a small reduction in binding to statistically significant impairments (Fig. 3D, TNF and IκBα promoters). In contrast, binding of RelA to the IFNβ promoter was not affected (Fig. 3D, right). Recruitment of RNA PolII to promoters of NF-κB target genes, including IκBα, was also compromised by Epi, and in a more pronounced and statistically significant way, by Acla (Fig. 3E). These results are consistent with previous observations that Doxo treatment strongly reduced the association between RelA and DNA, which was proposed a consequence of defective post-transcriptional modifications in NF-κB subunits (Ho et al., 2005). Furthermore, Doxo is also responsible for the reduced recruitment of at least one other transcription factor, HIF-1α, to its targets (Tanaka et al., 2012).

RelA consists of a REL-homology domain (RHD) and a transactivation domain (TAD, Supplementary Fig. 4D). The RHD includes the DNA binding domain (DBD), the dimerization domain (DD) and the nuclear localization sequence (NLS). DBD-mediated binding of RelA to regulatory regions of target genes dictates much of the diversity of NF-κB transcriptional responses (Toledano et al., 1993). We selected Epi to assess which RelA domain is targeted. Two different chimeric DNA constructs were tested: i) RelA RHD fused to a TAD derived from the Herpes simplex virus VP16 protein (RelA (2-320)-VP16); and ii) DBD from the bacterial tetracycline repressor (TET) fused to the RelA TAD (TET-RelA (286-551)) (Anrather et al., 1999). Similarly to the results obtained with full length RelA (Fig. 2F), Epi significantly inhibited the transcriptional activity of RelA (2-320)-VP16, as quantified in HEK293 cells transiently co-transfected with the κB luciferase reporter (Fig. 3F). However, Epi failed to inhibit the transcriptional activity of both TET-RelA (286-551) and TET fused to VP16 (TET-VP16) construct, as observed using a tetracycline operon luciferase (tet°-luc) reporter (Fig. 3F). This suggests that Epi decreases RelA transcriptional activity by targeting the RHD. Since Epi does not affect the TAD, this result suggests a mechanism of action different from NF-κB regulation by Doxo and Dauno in cancer cells (Campbell et al., 2004).

### Epirubicin and Aclarubicin bind to a κB-33 promoter sequence

To understand how anthracyclines affect binding of NF-κB to its targets, we used NMR and biophysical experiments. We tested a 14-mer DNA sequence (5’-CTGGAAATTTCCAG-3’) derived from the NF-κB-33 promoter (Chen et al., 1998, 2000). As expected, we detected binding of Epi and Acla to the DNA duplex by NMR spectroscopy, with dramatic changes in the imino region of the DNA after the addition of the drugs (Fig. 4A, B (black and red spectra), region 11-14 ppm) suggesting that the anthracyclines disturb the Watson-Crick hydrogen bonds in the double helix. Using isothermal titration calorimetry (ITC), we determined that two molecules of Epi or Acla bind to the DNA duplex with equilibrium dissociation constants (*K*_D_) of 11.4 μM and 11.7 μM, respectively (Table 1 and Supplementary Fig. 5A, B), indicating that Acla and Epi bind to the κB-33-derived DNA sequence with similar affinity. The affinity of Acla to different DNA molecules has been reported to be in the low nM to low µM range (Furusawa et al., 2016; Skovsgaard, 1987; Utsuno and Tsuboi, 1997), whilst Doxo, which differs from Epi in the stereochemistry of the 4’-OH group, binds to DNA in the nM range (Katenkamp et al., 1983). Most of the anthracyclines have a preference for GC-rich or TG-rich sequences, with the aglycon chromophore of the anthracyclines intercalating at a pyrimidine–purine step, and the sugar part interacting with the DNA minor groove (Chaires, 2015; Frederick et al., 1990; Temperini et al., 2003). The NF-κB-33 DNA oligonucleotide used in our assays contained two 5’-TG-3’ motifs suggesting that two molecules of anthracyclines can bind to it. When we used size exclusion chromatography (SEC) coupled with static light scattering (SLS) of RelA incubated with DNA and excess anthracyclines, an increase in the molecular weight (MW) of the DNA was observed, further pointing to binding of anthracyclines to DNA. However, due to their small MW, it was difficult to establish an accurate stoichiometry, and ITC proved to be more reliable, as discussed above (Supplementary Fig. 6, Table S2).

**Figure 4.**
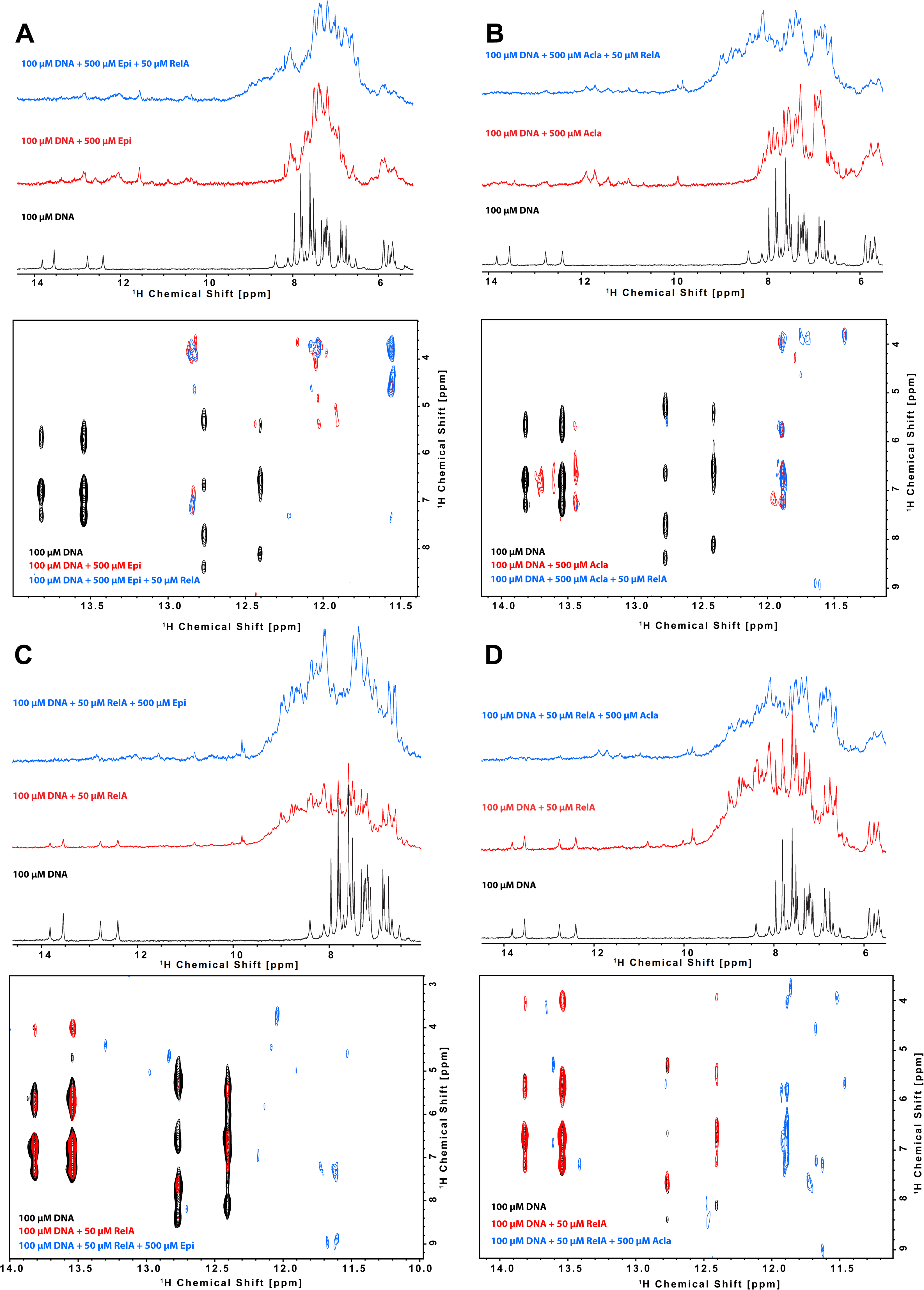
Binding of anthracyclines to *κ*B-33 promoter DNA and their effect on RelA-DNA complex following the DNA imino protons by NMR. 1D ^1^H spectrum (top) and zoom of the cross peaks of the imino protons with the deoxyribose in a 2D ^1^H, ^1^H-NOESY (bottom) of a 100μΜ 14*-*mer duplex DNA solution and subsequent addition of **A**, 500μΜ Epirubicin (Epi, red) and 50μΜ ^2^H, ^15^N-RelA_19-291_ dimer (blue), **B**, 500μΜ Aclarubicin (Acla, red) and 50 μΜ ^2^H, ^15^N-RelA_19-291_ dimer (blue), **C**, 50μΜ ^2^H, ^15^N-RelA_19-291_ dimer (red) and 500μΜ Epi (blue) and **D**, ^2^H, ^15^N-RelA_19-291_ dimer _1_ (red) and 500μΜ Acla (blue). Experiments were recorded at 800MHz and 10 °C in 100mM d11-Tris-HCl pH 7.5, 75mM NaCl, 5mM d10-DTT and 10% D_2_O.

**Table 1.** Summary of the data obtained by Isothermal Titration Calorimetry (ITC, data shown in Supplementary Figure 5). The ligand (or titrant) is titrated to the analyte (or titrand) in the cell, e.g. Epi to DNA. The measured thermodynamic properties are: N the stoichiometry of the titrant, KD the dissociation constant, ΔH enthalpy change, and ΔS the entropy change; T is the measurement temperature.

| | $K_D$ ($\mu\text{M}$) | N | $\Delta H$ (kJ/mol) | $-\text{T}\Delta S$ (kJ/mol) |
| --- | --- | --- | --- | --- |
| Epi to DNA | $11.3 \pm 4.6$ | $1.83 \pm 0.14$ | $-14.1 \pm 2.2$ | -14.2 |
| Acla to DNA | $11.7 \pm 2.0$ | $2.13 \pm 0.07$ | $-38.4 \pm 2.0$ | 10.3 |
| DNA to RelA | $0.23 \pm 0.03$ | $0.90 \pm 0.01$ | $40.7 \pm 0.9$ | -78.6 |
| (DNA + Epi) to RelA | $1.9 \pm 0.17$ | $0.88 \pm 0.04$ | $59.4 \pm 3.0$ | -92.1 |
| DNA to (RelA + Epi) | $0.98 \pm 0.12$ | $0.81 \pm 0.01$ | $-305 \pm 9$ | 271 |
| DNA to (RelA + Acla) | $0.74 \pm 0.32$ | $0.91 \pm 0.07$ | $-335 \pm 40$ | 300 |
| Epi to RelA | ND. Too weak |  |  |  |
| Acla to RelA | ND. Too weak |  |  |  |

### RelA RHD binds tightly to NF-κB-33

Regarding NF-κB binding to its target promoters, the published crystal structure of RelA RHD in complex with a 18-mer DNA segment derived from the κB-33 promoter revealed that the double-stranded DNA mainly contacts the interface between the DD and the DBD, with one of the protein monomers recognizing the 5’-GGAA-3’ site, and the other the 5’-GAAA-3’ site (Supplementary Fig. 7A) (Chen et al., 1998). Using NMR spectroscopy, we confirmed binding of the 14-mer DNA to RelA RHD containing the DD and DBD domains (residues 19-291, Fig. 4C, D (black and red spectra), Supplementary Fig. 7B). As expected, the DNA thymine and guanine imino’s remain detectable upon RelA addition but undergo line broadening consistent with the DNA maintaining its double helix structure upon protein binding (Fig. 4C, D (black and red spectra)); at the same time, addition of DNA to RelA causes chemical shift perturbations and line broadening of backbone amides on the DNA binding site (Supplementary Fig. 7). In addition, titration of DNA to RelA by ITC revealed strong binding, with an affinity of 230 nM (Table 1 and Supplementary Fig. 5C). This value is very similar to the reported value of 256 nM obtained using fluorescence polarization with a murine RHD construct and a 18-mer DNA using fluorescence polarization (Chen et al., 2000). Notably, the DNA-protein interaction is endothermic, i.e. enthalpically unfavorable and thus entropy-driven. By combining SEC and SLS experiments, we confirmed that both the protein and the DNA form dimers (Supplementary Fig. 6A, B and Table S2) but protein-DNA complexes were not observed, possibly due to a very fast off-rate (Bergqvist et al., 2009).

### Epirubicin and Aclarubicin disturb RelA-κB-33 binding

We then studied the effect of Epi and Acla on RelA-DNA binding by NMR, by following the effect either on the DNA or on the protein. Binding of RelA to the DNA did not substantially affect the DNA double helix as judged by DNA imino protons in thymine and guanine bases, as their intensity and chemical shifts were not substantially changed (Fig. 4C, D; black and red spectra). However, subsequent addition of anthracyclines clearly altered the Watson Crick base-paring even when bound to RelA (Fig. 4C, D; red and blue spectra). In contrast, titration of DNA with anthracyclines caused severe line broadening of the DNA base imino groups, which remained altered upon RelA addition (Fig. 4A, B). As the anthracyclines are expected to bind to the TG base pairs in the κB sequence located at the ends of the double helix (Fig. 5A), the DNA helix - though distorted - is still able to bind to RelA.

**Figure 5.**
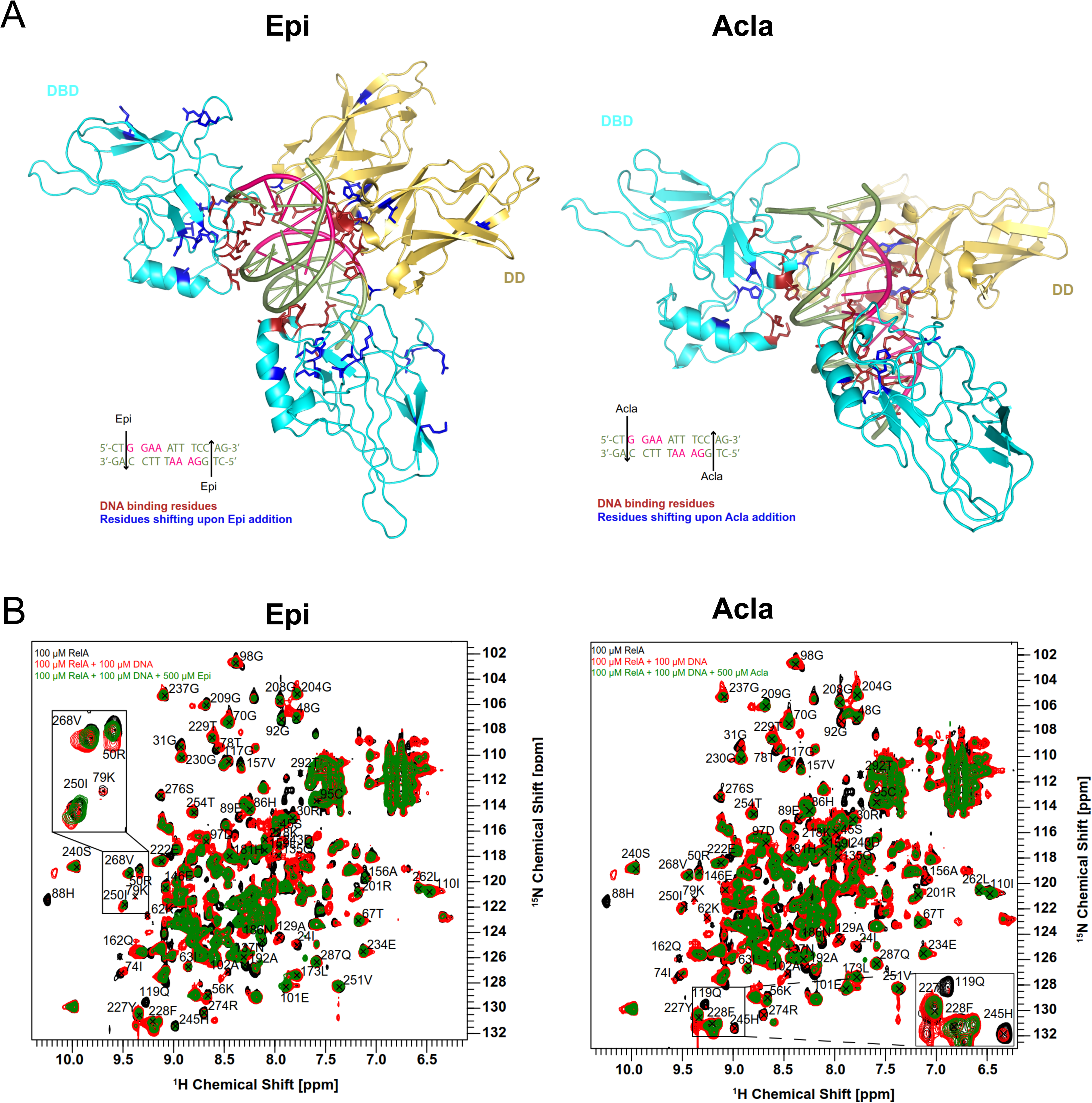
Epirubicin (Epi) and Aclarubicin (Acla) disturb RelA-DNA binding. **A,** Crystal structure of RelA REL-homology domain (RHD), containing the DNA binding domain (DBD) and the dimerization domain (DD), in complex with *κ*B-33 promoter DNA target (PDB code 1RAM). The RelA DBD and DD are represented as cartoon in cyan and yellow, respectively, whereas the DNA in olive green and pink. DNA highlighted in pink corresponds to the two half sites recognized by each RelA monomer. DNA binding residues are depicted in red, whereas the amino acid residues that shift after addition of Epi (**A**, left) and Acla (**A**, right) are depicted in blue; **B**, Superposition of 2D ^1^H,^15^N NMR correlation spectra of 100μΜ ^2^H, ^15^N-RelA_19-291_ free dimer (black), and in the presence of 100μΜ κB-33 14-mer DNA duplex (red) and upon additional presence of 500μΜ Epi (green, left) or 500μΜ Acla (green, right). All spectra were recorded at 800MHz, 20 °C, 16 scans.

Protein-based titration experiments to determine how RelA binding to DNA is affected by anthracyclines were performed in two ways: 1, RelA was first titrated with DNA and then with anthracyclines and 2, RelA was initially titrated with anthracyclines and then with DNA. Addition of anthracyclines to RelA titrated with DNA caused chemical shift perturbations and line-broadening beyond detection of some backbone resonances consistent with Epi and Acla perturbing the RelA-DNA complex but not disrupting it (Fig. 5B, compare green and red spectra) as the final spectra (green) do not resemble free RelA spectra (black). The RelA residues affected by the addition of the anthracyclines largely coincide with the DNA binding region. Amino acid residues in RelA-DNA complex particularly affected by Epi were 50R, 56K, 62K, 78T, 79K, 88H, 89E, 119Q, 129A and 157V in the DBD and 192A, 245H, 250I and 268V in the DD (Fig. 5A, B (left)). Addition of Acla to RelA-DNA complexes caused precipitation of the protein, which complicated any measurements. Yet, residues 88H, 119Q, 129A and 186N in the DBD and 245H in the DD could be analyzed and were altered (Fig. 5A, B (right)). Nevertheless, Epi affected the RelA-DNA complex more than Acla (Fig. 5B). Furthermore, addition of Epi to the RelA-DNA complex caused NMR chemical shift changes for some backbone amide signals in RelA (268V, 250I and 50R) towards an unbound state (Fig. 5B left zoomed-in), an effect that was not observed upon Acla addition (Fig. 5B right). Because the final spectra are not identical (compare green spectra in Fig. 5B), Epi and Acla should have different effects on the RelA-DNA complex. Differences in binding likely reflect the distinct chemical moieties involved, particularly because Acla contains three sugar rings while Epi contains one. In fact, studies showed that the sugar moieties of the anthracyclines bind to the minor groove of the DNA (Temperini et al., 2003, Chaires, 2015, Frederick et al., 1990). This may be compatible with the 3D structure of RelA-κB-33 (PDB code 1RAM, Fig. 5A, Supplementary Fig 7A), with the TG sequences at the end of the DNA duplex and with the accessible minor groove being able to accommodate the anthracycline sugars and in close vicinity of DD. Indeed, our results suggest that Epi and Acla bind to RelA through their sugar parts (Supplementary Fig. 8). Notably, addition of Epi to RelA followed by addition of DNA led to different spectra (Supplementary Fig. 9A, compare red and green spectra), suggesting that the final complexes are slightly different. This is more complex for Acla spectra (Supplementary Fig. 9B, compare red and green spectra), which is not conclusive because addition of Acla to RelA leads to protein precipitation, and the final spectrum obtained after DNA addition (green) is therefore much weaker than its counterpart (red).

To further explore how anthracyclines affect the RelA-DNA complex we performed ITC competition experiments. The binding affinities of DNA to a pre-formed complex of RelA with Epi (980 nM) or Acla (740 nM) were weaker than the binding affinities of the DNA to the protein alone (230 nM) (Table 1). Moreover, the thermodynamic features of the interaction were different: binding of DNA to RelA is entropy-driven and endothermic (positive ΔH), whilst binding of DNA to RelA incubated with anthracyclines is enthalpy-driven and exothermic (negative ΔH). Surprisingly, when DNA with a 2-fold excess of Epi was titrated to the protein, the binding affinity decreased dramatically from a *K*_D_ of 230 nM to 1900 nM but the reaction is similarly endothermic (Table 1). This suggests that when the anthracyclines-DNA complex is formed, binding to RelA is mainly affected.

We also tested a potential direct binding between anthracyclines and RelA. Whereas no binding was detected by ITC, likely because this method is not suitable for very weak interactions (Table 1), NMR spectroscopy showed that both Epi and Acla bind weakly to the protein (Supplementary Figs. 8, 10). We did not observe significant RelA backbone amide changes after the addition of anthracyclines in 2D-TROSY NMR protein spectra, indicating that Epi and Acla bind very weakly to the protein (Supplementary Fig. 10). Binding to RelA, though weak, is mediated through the sugar part of the anthracyclines, as observed in saturation transfer difference (STD) experiments (Supplementary Fig. 8A,B). In agreement, Doxo-none, which lacks the sugar moiety, did not bind to RelA (Supplementary Fig. 8C).

## Discussion

We show that anthracyclines inhibit NF-κB-dependent gene expression of critical pro-inflammatory mediators such as TNF in macrophages (Fig. 1A, 2A) while not all cellular transcription was diminished (Fig. 2D). Despite the general notion that anthracyclines induce NF-κB-dependent anti-apoptotic gene expression when used as chemotherapeutic drugs (Wang et al., 1996) (Arlt et al., 2001) (Janssens et al., 2005) (Van Antwerp et al., 1996) (Wu et al., 1996), Doxo and Dauno have also been described as repressors of NF-κB activity (Campbell et al., 2004) (Ho et al., 2005). It is highly likely that the final outcome of anthracycline treatment is dependent on cell and tumor type. Using transcriptomics, we found that only a very small proportion of inflammation-driven genes were upregulated by either Epi or Acla (Fig. 2A) while none of the tested anti-apoptotic factors, including cellular inhibitor of apoptosis proteins (cIAPs), were affected. Instead, we detected downregulation of a broad range of pro-inflammatory genes in Epi- or Acla-treated macrophages, that all have NF-κB as the common regulator of their transcription. How anthracyclines inhibit transcription by NF-κB can be gene-dependent, as multiple factors are known to dictate transcriptional activity from κB sites, including the composition of the bound NF-κB dimers, interactions amongst NF-κB subunits at tandem binding sites, as well as regulation by other transcription factors (Mulero et al., 2017). However, similar ChIP results at several NF-κB target promoters in combination with the Epi- and Acla-dependent decreased binding of RelA and PolII upon pro-inflammatory stimulation, exposed a novel mechanism mediated by anthracyclines that implied a direct role in counteracting NF-κB activity at its binding sites (Fig 3D and E).

Because Epi is protective in sepsis through ATM-mediated signaling (Figueiredo et al., 2013), and due to the genotoxicity of this class of drugs, we wondered whether cytokine regulation by anthracyclines depended on the induction of DNA damage responses (DDRs). We used a selection of anthracyclines that, unlike Epi, do not cause detectable DNA damage in the conditions tested (Qiao et al., 2020). We used the anthracyclines Acla and diMe-Doxo that do not induce DNA damage (but still allowed histone release at defined sides in the genome) to test whether DNA damage was critical in this process. Not only was pro-inflammatory gene expression orchestrated by NF-κB still downregulated by Acla, Doxo-none and the newly developed diMe-Doxo (Fig. 1F, 1H), but also the DDR was not required for cytokine suppression by using an ATM inhibitor and *Atm^-/-^* BMDMs (Fig. 1A, 1B and Supplementary Fig. 1A). In agreement with this, DNA damaging drugs such as Eto did not modulate cytokine production, highlighting the lack of interdependency between regulation of NF-κB-dependent transcription and DNA damage. Whereas the anticancer activity of the anthracyclines has been attributed to stabilization of the TopoII-DNA cleavable complex and consequent formation of DNA double-strand breaks, anthracyclines lacking DNA damaging competence are also effective, such as Acla for the treatment of acute myeloid leukemia (AML)(Qiao et al., 2020). Previously unrecognized properties of the anthracyclines are now known to decisively contribute to cancer treatment, as is the case of chromatin damage caused by histone eviction (Pang et al., 2013; Yang et al., 2013), and thus it is likely that regulation of NF-κB activity also proves to be important for the therapeutic efficacy. Limiting inflammation through NF-κB modulation may have been an overlooked aspect of Epi, Doxo and Acla with a fundamental role in the clinical success of this class of drugs. Future therapeutic strategies are expected to focus on anthracyclines without DNA damaging activities, in an attempt to overcome the severe side effects of these drugs. Indeed, diMe-Doxo and Acla -both lacking DNA damage activity- are not cardiotoxic and do not induce second tumors(Qiao et al., 2020) and could be more suitable for controlling sepsis and other inflammatory responses. Importantly, the novel role in NF-κB suppression will also be part of these treatment options as we showed that, although with different efficacies, it is common to all anthracyclines tested and independent of DNA damage, and low doses seem to be sufficient to prompt this regulatory effect.

But can we understand how anthracyclines control NF-κB? Using NMR we showed that Epi and Acla disturb the complexes formed between the NF-κB subunit RelA RHD and its cognate DNA κB-33 binding sites (Figs. 4, 5, Table 1). Epi and Acla bind to κB-33 DNA and distort its conformation, by probably intercalating between the TG bases at the ends of the κB-33 promoter and with the anthracycline sugars interacting with the DNA minor groove. The anthracyclines sugars also interact weakly with RelA, promoting the formation of a ternary complex between anthracyclines-DNA-RelA. In this ternary complex, the double helix complex with anthracyclines is still able to bind to RelA RHD, though with a lower affinity. Whereas the two anthracyclines were shown to perturb the binding of RelA to DNA differently, the results from NMR and biophysical experiments did not explain why Acla is the strongest regulator of NF-κB-dependent transcription. It is possible that the presence of the extra sugars in Acla is responsible for a stronger interaction with RelA RHD and the DNA, but that this ternary complex leads to a lower engagement of TAD, transcription factors and RNA polymerase for transcription. It will be important to structurally dissect which other effects contribute to the efficient Acla-mediated NF-κB transcriptional suppression and to explore promoter-specific effects. We observed that the RelA residues affected by the addition of Epi and Acla to a RHD-DNA complex are located in both the DBD and the DD of the RHD (Fig. 5). These binding results corroborate the reporter assays that revealed that Epi downregulates RelA transcriptional activity by targeting the RHD (Fig. 3F). Epi and Acla also bind with low affinity through their sugar parts to RelA, which was confirmed by comparing with Doxo-none that lacks the sugar and did not show any binding (Supplementary Fig. 8). The relevance of anthracycline-RelA binding for the transcriptional suppression is suggested by the observation that Doxo-none is a weaker modulator of TNF when compared with Epi, Doxo and Acla (Fig. 1F). However, diMe-Doxo, with a sugar moiety that resembles both Epi (contains only one sugar) and Acla (the amine group is dimethylated), is weaker than the classical anthracyclines at regulating the genes tested (Fig. 1H). We are currently investigating its RelA and DNA binding properties to uncover potential new rules affecting complex formation.

Epi and Acla have a different structure. Acla contains three sugar rings and Epi only one aminosugar. The sugar moieties of anthracyclines occupy the minor groove of DNA and the amino group at the 3′ position of the sugar are important for DNA binding affinity and antitumor activity (Temperini et al., 2003; Wander et al., 2020; Zunino et al., 2001). Both Epi and Acla were shown to interact with DNA bases within the κB motif, and our ITC competition experiments suggest that this interaction can disturb the RelA-DNA complex formation (Supplementary Fig. 5). Noticeably, the engagement of RelA and other NF-κB factors with κB sites is highly dynamic and regulated by chromatin organization and other nuclear factors.

We have dissected the contribution of anthracyclines in the control of NF-κB controlled inflammatory responses that can lead to sepsis. Using a series of dedicated variant drugs with defined activities, we could dissociate the DNA damaging from the histone eviction activities. We show that controlling inflammatory responses does not require DNA damage. In fact, using biophysical methods and NMR we show that anthracyclines disturb the interaction of the NF-κB subunit RelA in the promoter region thus suppressing the production of pro-inflammatory signals at very low concentrations. We thus uncover a new mechanism of action of the anthracycline class of anti-cancer drugs that can bridge application in situations of uncontrolled inflammatory responses, such as sepsis.

## STAR Methods

### Key Resources Table

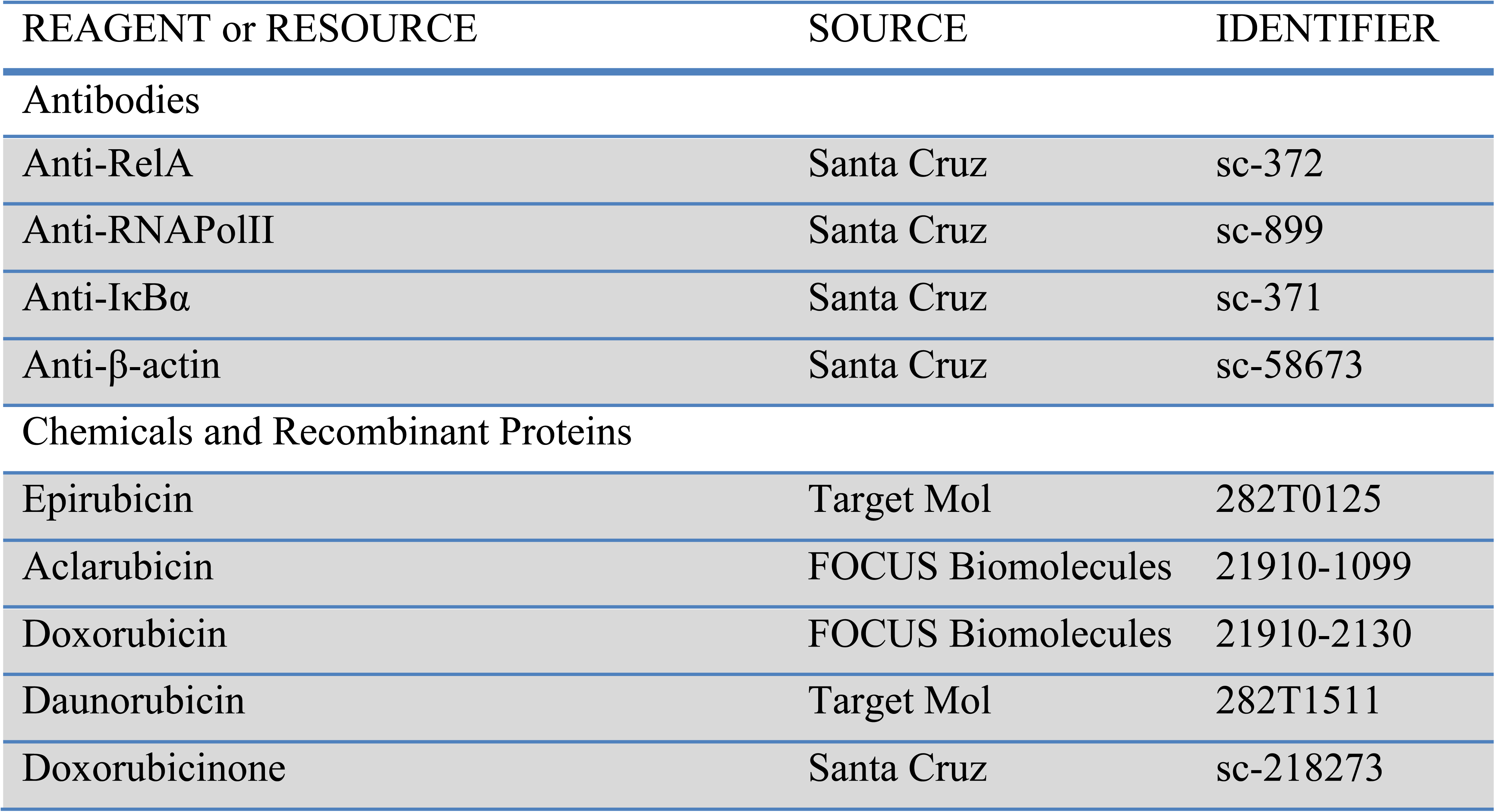
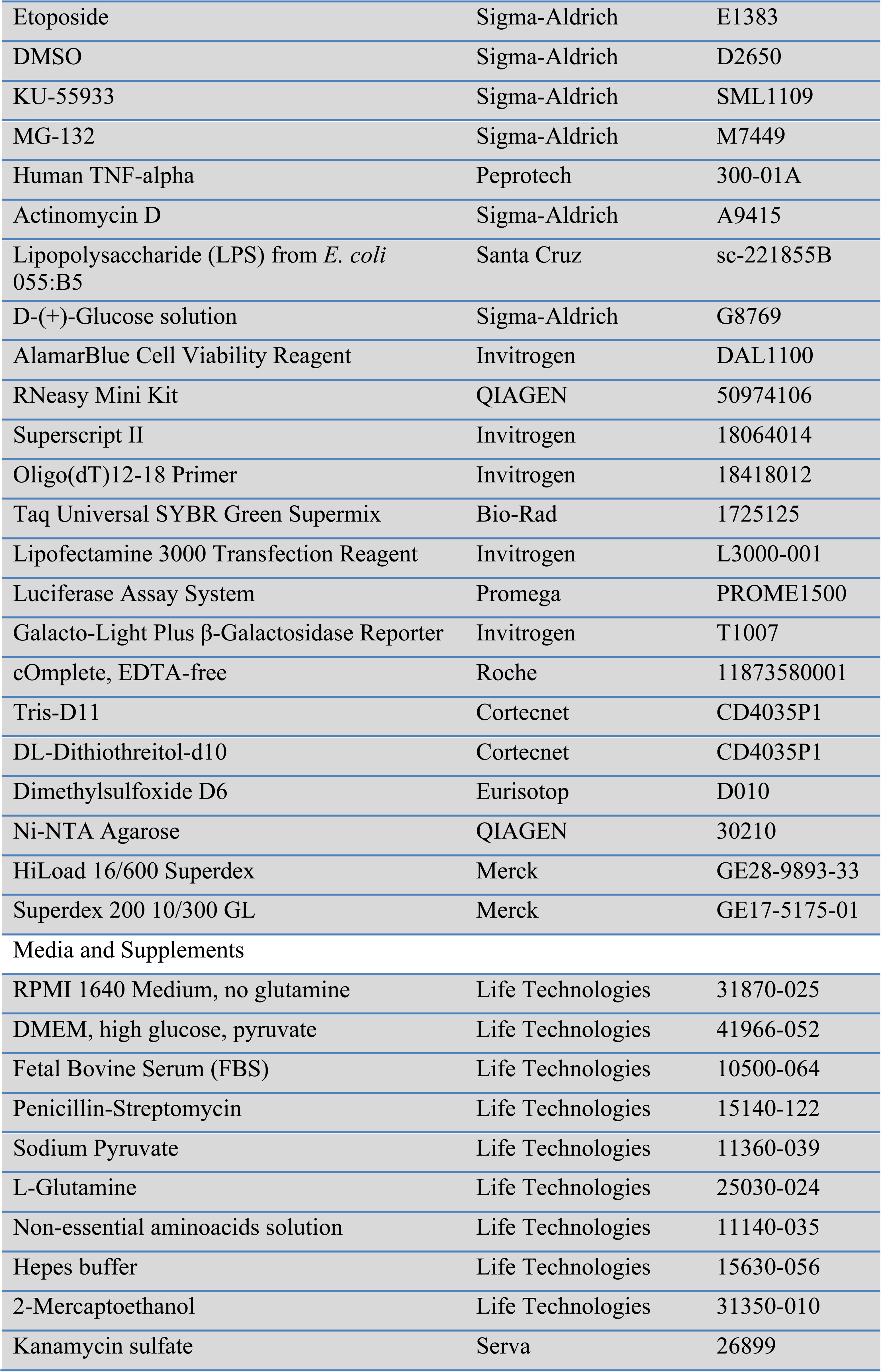
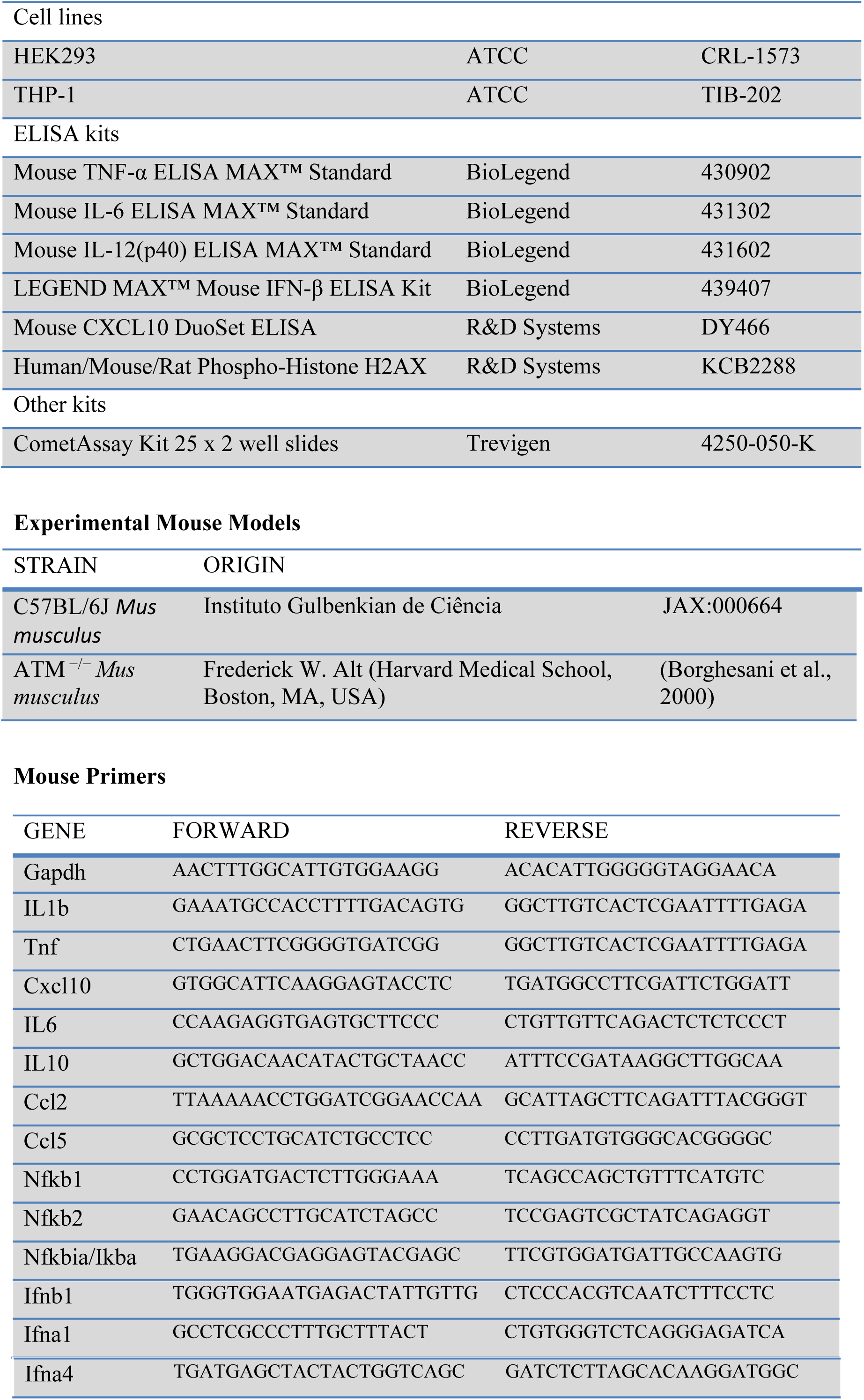

### Resource Availability

Further information and requests for resources and reagents should be directed to and will be fulfilled by the Lead Contacts: Ana Neves-Costa; and Luis F. Moita.

### Experimental Models

#### Mice

All animal studies were performed in accordance with Portuguese regulations and approved by the Instituto Gulbenkian de Ciência ethics committee and DGAV. Atm^−/−^ mice (Borghesani et al., 2000) and C57BL/6J control mice were bred and maintained under specific-pathogen free (SPF) conditions at the Instituto Gulbenkian de Ciência with 12 h light/12 h dark cycle, humidity 50%–60%, ambient temperature 22 ± 2°C and food and water ad libitum. Male mice of 8 to 12 weeks old were used in the experiments and age-matched mice were randomly assigned to experimental groups.

#### Primary cell cultures

For the generation of bone marrow-derived macrophages, total bone marrow cells were flushed from femurs and tibiae, counted and seeded (3×10^6^ cells/mL) in RPMI 1640 supplemented with 10% (v/v) FBS, 0.2% (v/v) Penicillin-Streptomycin, 1% (v/v) Sodium Pyruvate, 1% (v/v) L-Glutamine, 1% (v/v) non-essential aminoacids, 1% (v/v) Hepes buffer and 0.05mM of 2-Mercaptoethanol (all Gibco, Life Technologies) supplemented with 30% conditioned medium from mouse macrophage colony stimulating factor (M-CSF)-producing L929 cells. On day 7, adherent BMDM cells were collected, counted, reseeded in 96-well plates (5×10^4^/well) and treated and/or challenged as indicated. HEK293 cells were obtained from ATCC and cultured in DMEM supplemented with 10% (v/v) FBS and 1% (v/v) Penicillin-Streptomycin (all Gibco, Life Technologies). Cells were cultured in a humidified atmosphere containing 5% CO_2_.

### Method details

#### Reagents

Epirubicin, Aclarubicin, Doxorubicin, Daunorubicin, Doxorubicinone and Dimethyl-doxorubicin were dissolved in PBS at 1mg/mL, stored at -80°C and used at the indicated final concentrations. Anthracyclines were from various commercial sources except for Dimethyl-doxorubicin, a gift from Jacques Nefjees. Etoposide (Sigma-Aldrich) was dissolved in DMSO and used at indicated final concentrations. MG-132 (Sigma-Aldrich) was dissolved in DMSO and used at a final concentration of 10µM. ATM inhibitor KU-55933 (Sigma-Aldrich) was dissolved in DMSO and used at a final concentration of 5µM. Human recombinant TNF (PeproTech) was dissolved in RPMI and used at a final concentration of 10ng/mL for the indicated times. Actinomycin D (ActD, Sigma-Aldrich) was dissolved in DMSO and used at a final concentration of 5μg/mL for the indicated times. LPS (from *E. coli* 055:B5, Santa Cruz) was dissolved in RPMI and used at a final concentration of 100ng/mL. PFA-fixed *E. coli* were prepared as before (Moura-Alves et al., 2011) in house and added to the medium at a ratio of 20 bacteria per cell. AlamarBlue cell viability assay (Invitrogen) was used to determine cell viability according to the manufacturer’s instructions.

#### E. coli-induced sepsis model

This model has been described in detail before (Colaço et al., 2021). Briefly, a starter culture from a single *E. coli* colony was grown overnight at 37°C with agitation in Luria-Bertani broth (LB). The following morning, the culture was diluted 1:50 in LB and incubated for 2.5 h until late exponential phase was reached (OD600nm = 0.8-1.0). The culture was then centrifuged at 4400g for 5 min at room temperature, washed and ressuspended in PBS to OD600nm = 4.5-5.0, corresponding to 1-2×10^9^ CFU/mL. This bacterial suspension was immediately injected intraperitoneally (200μL/mouse) in mice using a 27G-needle, always in the morning. Epirubicin or Aclarubicin were dissolved in PBS and injected intraperitoneally (200μL/mouse) at 0.5 and 0.73μg/g body weight, respectively, at the time of infection. Mice were sacrificed at 8h by CO2 inhalation and blood was collected by cardiac puncture. Serum was collected after centrifuging the blood at 1600g for 5 min.

#### Cytokine production measurement

Mouse sera and cell culture supernatants were collected at indicated time points and TNF, Cxcl10, IL6, IL-12p40 and IFNβ production quantified using ELISA kits (BioLegend and R&D Systems) according to the manufacturer’s instructions.

#### qRT-PCR

Total RNA was isolated from BMDM using the RNeasy Mini Kit (QIAGEN), reverse transcribed with Superscript II reverse transcriptase (Invitrogen) using oligo(dT)12-18 primers. Specific RNA specimens were quantified by PCR reaction using SYBRgreen Supermix (Bio-Rad) on the QuantStudio7 Flex real-time PCR system (Applied Biosystems). Cycle thresholds (Ct), normalized to Ct of Gapdh, were used to calculate fold increase over control. Primer sequences are described in the supplementary information section.

#### RNA-seq and data analysis

Total RNA was extracted as described above and quality was assessed using the AATI Fragment Analyzer. Samples with RNA Quality Number (RQN) >8 and clearly defined 28S and 18S peaks were further used for preparation of mRNA libraries, which were pooled and sequenced (75bp, single end) using NextSeq500. The quality of the sequences was assessed using FASTQC and MultiQC before the alignment (Ewels et al., 2016). Sequences were aligned against the *Mus musculus* genome version 89, with the annotation file for the genome version 89, both from Ensembl. The alignment was done with STAR (Dobin et al., 2013), using the default parameters and including the GeneCounts option. The files from GeneCounts were imported to R (version 3.5.1), taking into account the strandness associated with the sequencing protocol. DESeq2 (version 1.22.1) (Love et al., 2014) was used for the downstream analysis. Heatmaps were created with data normalized from raw counts through Regularized Log Transformation (rlog) (Love et al., 2014). The log2FC was shrunk using the ‘ashr’ (Adaptive SHrinkage) package (Stephens, 2017) and genes were considered differentially expressed when the p-value was below 0.05 after adjusting using false discovery rate (FDR). Gene Information was obtained from org.Mm.eg.db. Functional clustering was performed using the DAVID Gene Functional Classification Tool (https://david.ncifcrf.gov).

#### Antibodies used in Western Blot and Immunofluorescence

The following antibodies were used for the specific detection of: IκBα, sc-371, Santa Cruz; RelA, sc-372, Santa Cruz; and β-actin, sc-58673, Santa Cruz. Primary antibodies were detected using HRP-conjugated secondary antibodies (Cell Signaling).

#### Transient transfection and Reporter assay

N-terminal Myc-tagged RelA, RelA (2– 320)/VP16, TET/RelA (268–551) and TET/VP16 expression vectors, the NF-κΒ firefly luciferase reporter construct (*κB-luc*), and the tetracycline operon (tet°) firefly luciferase reporter (*tet°-luc*), were previously described (Anrather et al., 1999). The κB sequences in the *κB-luc* luciferase reporter are: 5’-TGCTGGGAAACTTTC-3’ and 5‘-TGCTGGGAATTCCTC-3′. The pSV-β-galactosidase reporter consists of the lacZ gene from *E. coli* under the control of the SV40 early promoter and enhancer. Transient transfections were carried out using the lipofectamine 3000 transfection reagent (Invitrogen). 24 hours after transfection, cells were either pre-exposed to Epirubicin for 1 hour prior to TNF stimulation or treated with Epirubicin for 16 hours in the RelA over-expression assays. After incubation, cells were lysed and firefly luciferase and β-galactosidase activity were measured using the Luciferase Assay System (Promega) and Galacto-Ligh System (Invitrogen), respectively, following the manufacturer’s instructions.

#### Chromatin Immunoprecipitation (ChIP)

Chromatin immunoprecipitation (ChIP) was performed on HEK293 cells as previously described (De Almeida et al., 2011). Antibodies against RelA (sc-372, Santa Cruz) and RNAPolII (sc-899, Santa Cruz) were used for immunoprecipitation.

#### Comet assay

Comet assay was performed in THP-1 cells (ATCC catalog number TIB-202) using the CometAssay Kit 25 x 2 well slides (Trevigen catalog number 4250-050-K). A step-by-step protocol detailing the exact procedures and all the materials used is available online from Protocol Exchange (Alkaline Comet Assay using the monocytic cell line THP-1, https://doi.org/10.21203/rs.2.11936/v2).

#### phosphoH2AX quantification

Phosphorylation of histone H2AX at serine 139 was quantified by cell-based ELISA using the kit Human/Mouse/Rat Phospho-Histone H2AX (R&D catalog number KCB2288) according to the manufacturer’s instructions.

#### Quantification and Statistical Analysis

Data are expressed as mean values ± standard deviation. Mann-Whitney test was used for pairwise comparisons and two-way ANOVA with Tukey test was used for multiple comparisons. Statistical analysis was performed with GraphPad Prism 6.0 (GraphPad Software). The number of subjects used in each experiment is defined in figure legends. The following symbols were used in figures to indicate statistical significance: p < 0.05 (*); p < 0.01 (**); p < 0.001 (***); p < 0.0001 (****).

#### Purification of recombinant RHD RelA

A human RelA construct (residues 19-291) was prepared by subcloning into pETM11 vector, which contains an N-terminal His6-tag and a TEV protease cleavage site. The new construct contains the N-terminal DNA-binding domain (DBD, residues 19-191) and the C-terminal dimerization domain (DD residues 192-291). The plasmids were transformed into *E. coli* strain BL21 (DE3) cells and cultured overnight at 20°C in LB media supplemented with 100µg/mL kanamycin.

For the preparation of uniformly-labeled ^2^H (∼100%) RHD RelA, ^15^N (99%)-labeled protein was expressed at 37°C using M9 minimal medium containing ^15^NH_4_Cl, [^12^C]D-d7-glucose(2g/L) (97% *D*, Sigma-Aldrich) in 100% D_2_O. A standard protocol of sequential pre-cultures for better D_2_O adaptation over a 3d period was followed to increase the yield of protein expression in 100% D_2_O. On the first day, a 25mL preculture in LB medium was prepared and grown overnight at 37°C. The following day, a preculture of 50mL M9 minimal medium in H_2_O was inoculated with 1ml of the overnight LB preculture and grown at 37°C. After some hours, when the preculture reached an optical density at 600nm (OD_600_) close to 0.6, it was spun down for 10min at 3,202*g*. The cells were ressuspended in 1mL of M9 medium in 100% D_2_O and used for the inoculation of 100mL of M9 medium in 100% D_2_O, such that the OD_600_ was 0.1-0.15. This small culture was left overnight at 37°C. The next day, this culture was added to 900mL of M9 medium in 100% D_2_O. All cultures in minimal media were induced at OD_600_ of 0.8 with 0.5mM of IPTG overnight at 20°C.

After overnight induction, cell pellets were lysed by sonication in lysis buffer (50mM Tris-HCl pH 8, 300mM NaCl, 5mM imidazole, 5mM mercaptoethanol, 0.025mg/mL DNAse I, 0.1mg/mL Lysozyme, 2.5mM MgSO_4_, 0.1% NP-40 and 1 pill of protease inhibitor EDTA-free (cOmplete Tablets, Mini EDTA free, Roche) per 30mL lysate. The cell lysate was centrifuged at 60,000 *g* for 30min at 4°C. After filtration, the His-tagged protein in the supernatant was loaded on an IMAC (Immobilized Metal Affinity Chromatography). The supernatant was applied to Ni-NTA resin (QIAGEN) previously equilibrated with 3 column volumes of buffer A (50mM Tris-HCl pH 8, 300mM NaCl, 5mM imidazole, 5mM mercaptoethanol). Bound protein was washed with 3 column volumes of buffer A and unspecific bound protein was washed away with 3 column volumes of Wash Buffer (50mM Tris-HCl pH 8, 1M NaCl, 5mM Imidazole and 5mM mercaptoethanol). His_6_-tagged protein was eluted using elution buffer (50mM Tris-HCl pH 8, 300mM NaCl, 300mM Imidazole and 5mM mercaptoethanol). The affinity His-tag was removed from the protein by TEV (1:5 protein:TEV ratio) cleavage during dialysis into 50mM Tris-HCl pH 8, 300mM NaCl and 5mM mercaptoethanol buffer overnight at 4°C. The cleaved tag and TEV protease were removed from the target protein using a second IMAC step in dialysis buffer. The fractions containing RelA were pooled, concentrated and further purified by size-exclusion chromatography (SEC) using a Superdex 75 Hiload 16/60 column (S75, GE Healthcare, Merck). The SEC buffer used was 50mM Tris-HCl pH 7.5, 150mM NaCl, 1mM EDTA and 5mM Dithiothreitol (DTT). For the ^2^H, ^15^N-labeled RelA, prior to the SEC purification step, 2M urea were added to the protein sample for 1 hour, in order to enhance the proton chemical exchange. The final yields were 12.5mg for ^2^H, ^15^N RelA and 52mg for unlabeled RelA per liter of cell culture.

For NMR experiments, all protein samples were exchanged by successive concentration/dilution steps into NMR buffer (100mM d11-Tris-HCl (Cortecnet) pH 7.5, 75mM NaCl, and 5mM d10-DTT (Cortecnet), 90% H_2_O/10% D_2_O). The protein concentrations were calculated using the absorption at 280nm wavelength by using molar extinction coefficients of 17420 M^-1^ cm^-1^ for RelA_19-291_.

#### NMR Spectroscopy

One-dimensional (1D) ^1^H NMR experiments were recorded using a WATERGATE pulse sequence at 25°C on a Bruker AvanceIII 800MHz spectrometer equipped with a cryogenic TCI-probehead (^1^H, ^31^P, ^13^C, ^15^N) with Z-gradients. 1D ^1^H experiments were performed using a WATERGATE pulse sequence with 32k time domain and 128 scans in 100mM d11-Tris-HCl pH 7.5, 75mM NaCl, 5mM d10-DTT and 10% D_2_O. STD experiments were recorded using an interleaved pulse program with on-resonance protein irradiation at 0.15ppm for Epirubicin, 0.5ppm for Aclarubicin and 0.6ppm for Doxorubicinone and off-resonance irradiation at -5ppm with 2sec effective irradiation, using 800 scans and 32k time domain points (600MHz). Each experiment was performed using 500μM of compound 10μM of unlabeled protein. Reference STD experiments without protein were performed at the same conditions, using the same irradiation regions. Spectra were processed using TOPSPIN 3.2 (Bruker Biospin, Rheinstetten, Germany).

NMR binding studies were performed at 25°C using 100μM ^2^H(∼100%), ^15^N-labeled RelA_19-291_ dimer in a 100mM d11-Tris-HCl buffer (pH 7.5, 75mM NaCl, 5mM d10-DTT and 10% D_2_O) by adding compound to a final concentration of 500 and/or 100μM of duplex-DNA, and monitoring the changes by ^1^H, ^15^N TROSY experiments. A reference experiment was performed under the same conditions with the same volume of DMSO-d6 (Eurisotop) as used for the compound titration. Changes in the DNA structure were detected by recording NMR Imino NOESY spectra at 10°C, using 100μM duplex-DNA, to which compound (Epi and Acla) to a final concentration of 500 μM and 50 μM ^2^H(∼100%), ^15^N-labeled RelA_19-291_ dimer in a 100mM d11-Tris-HCl buffer (pH 7.5, 75mM NaCl, 5mM d10-DTT and 10% D_2_O) were added in a different order.

Chemical shift assignment of RelA_19-291_ was obtained at 950 MHz and 25°C using TROSY versions of 3D HNCACB, HNCA, HN(CO)CA, HN(CA)CO and HNCO experiments (Sattler et al., 1999),(Zhang et al., 1994) on a 200 μΜ ^2^H(∼100%), ^13^C, ^15^N-labeled RelA_19-291_ dimer in 100mM d11-Tris-HCl buffer (pH 7.5, 75 mM NaCl, 5 mM d10-DTT and 10% D_2_O) based on the assignment of three similar constructs of a ^15^N-labeled, perdeuterated RelA RHR (residues 19-325) in complex with perdeuterated p50 RHR (residues 37-363), a RelA DBD (residues 19-191) and a RelA DD (residues 190-321) in complex with deuterated p50 DD (residues 245-350) (Mukherjee et al., 2016). Assignment of RelA bound to DNA was performed by following the resonances during the DNA titration and confirmed by using a TROSY version of 3D HNCA on a 200 μΜ ^2^H(∼100%), ^13^C, ^15^N-labeled RelA_19-291_ dimer with 100μM duplex 14-mer DNA in 100mM d11-Tris-HCl buffer (pH 7.5, 75mM NaCl, 5mM d10-DTT and 10% D_2_O) obtained at 950MHz and 25°C. All datasets were processed using NMRPipe (Delaglio et al., 1995) and analyzed with CCPN analysis 2.4.2 (Vranken et al., 2005).

#### Static light scattering (SLS)

Static light scattering (SLS) experiments were performed at 303K using a Viscotek TDA 305 triple array detector (Malvern Instruments) downstream to an Äkta Purifier (GE Healthcare) equipped with an analytical size exclusion column (Superdex 75 or 200 10/300 GL, GE Healthcare, Merck) at 4°C. The samples were run in 50mM Tris-HCl pH 7.5, 150mM NaCl, 1mM EDTA and 5mM DTT with a concentration of 2mg/mL for the protein and at ratios of 1:2 protein:DNA and of 1:2:6 protein:DNA:anthracyclines at a flow rate of 0.5mL/min. The molecular masses of the samples were calculated from the refractive index and right-angle light-scattering signals using Omnisec (Malvern Instruments). The SLS detector was calibrated with a 4mg/ml BSA solution with 66.4kDa for the BSA monomer and a dn/dc value of 0.185mL/g for all protein samples.

#### Isothermal titration calorimetry

ITC measurements were carried out at 25°C using a MicroCal PEAQ-ITC (Malvern Instruments Ltd). The titrations were performed in 50mM HEPES pH 8.0, 100mM NaCl and 1mM mercaptoethanol and 1% DMSO. The calorimetric titration consisted of 19 injections of 1.5μL of a 125µM DNA sample, into the reaction cell containing 400μL of 25μM RelA or to 25μM RelA with 200μM Epirubicin/ Aclarubicin, at a stirring speed of 1000rpm. The heat of dilution was obtained by titrating DNA into the sample cell containing only buffer and this was subsequently subtracted from each experimental titration. For evaluating the effect of Epirubicin bound to DNA on binding to RelA, a calorimetric titration consisted of 19 injections of 1.5μL of a 125µM DNA with 250µM Epirubicin mixture, into the reaction cell containing 400μL of 25μM RelA. For the determination of the binding affinity of the compounds to the DNA, a calorimetric titration was performed consisting of 19 injections of 1.5μL of a 500µM compound sample, into the reaction cell containing 400μL of 50μM DNA. The heat of dilution was obtained by titrating compound into the sample cell containing only buffer and this was subsequently subtracted from each experimental titration. For the determination of the binding affinity of the DNA to the protein, a calorimetric titration was performed consisting of 19 injections of 1.5μL of a 125µM DNA sample, into the reaction cell containing 400μL of 25μM RelA. The ITC data were analyzed using the MICROCAL PEAQ-ITC analysis software provided by Malvern.

## Acknowledgements

We are grateful to the Genomics Unit and the Animal House at IGC. We thank Margarida Gama-Carvalho for the critical revision of the manuscript. This work was supported by the European Commission Horizon 2020 (ERC-2014-CoG 647888-iPROTECTION) and by Fundação para a Ciência e Tecnologia (FCT: PTDC/BIM-MEC/4665/2014). SW is funded by the German Research Foundation (4971/6-1), the German Federal Ministry for Education and Research (BMBF) (01EN2001) and by the Integrated Research and Treatment Center -Center for Sepsis Control and Care (CSCC) at the Jena University Hospital. The CSCC is funded by the German Ministry of Education and Research (BMBF No. 01EO1502).

## Author Contributions

Conceptualization, L.F.M. with input from A.N.-C.; Methodology and Formal Analysis: A.C., V.B., M.S., A.C.M. and A.N-C.; Investigation, A.C., D.P., E.K., N.P., H.C., R.G., A.B., K.W., T.V., C.F.M., I.S., P.P., S.C., F.B.M., A.G. and A.N.-C.; Resources, J.A.F., S.F.A., J.A., S.W., M.P.S., J.N; Original Draft, A.N.-C. and L.F.M.; Supervision A.N.-C. and L.F.M.; Funding Acquisition, L.F.M.

---

**Supplementary Figure 1.**
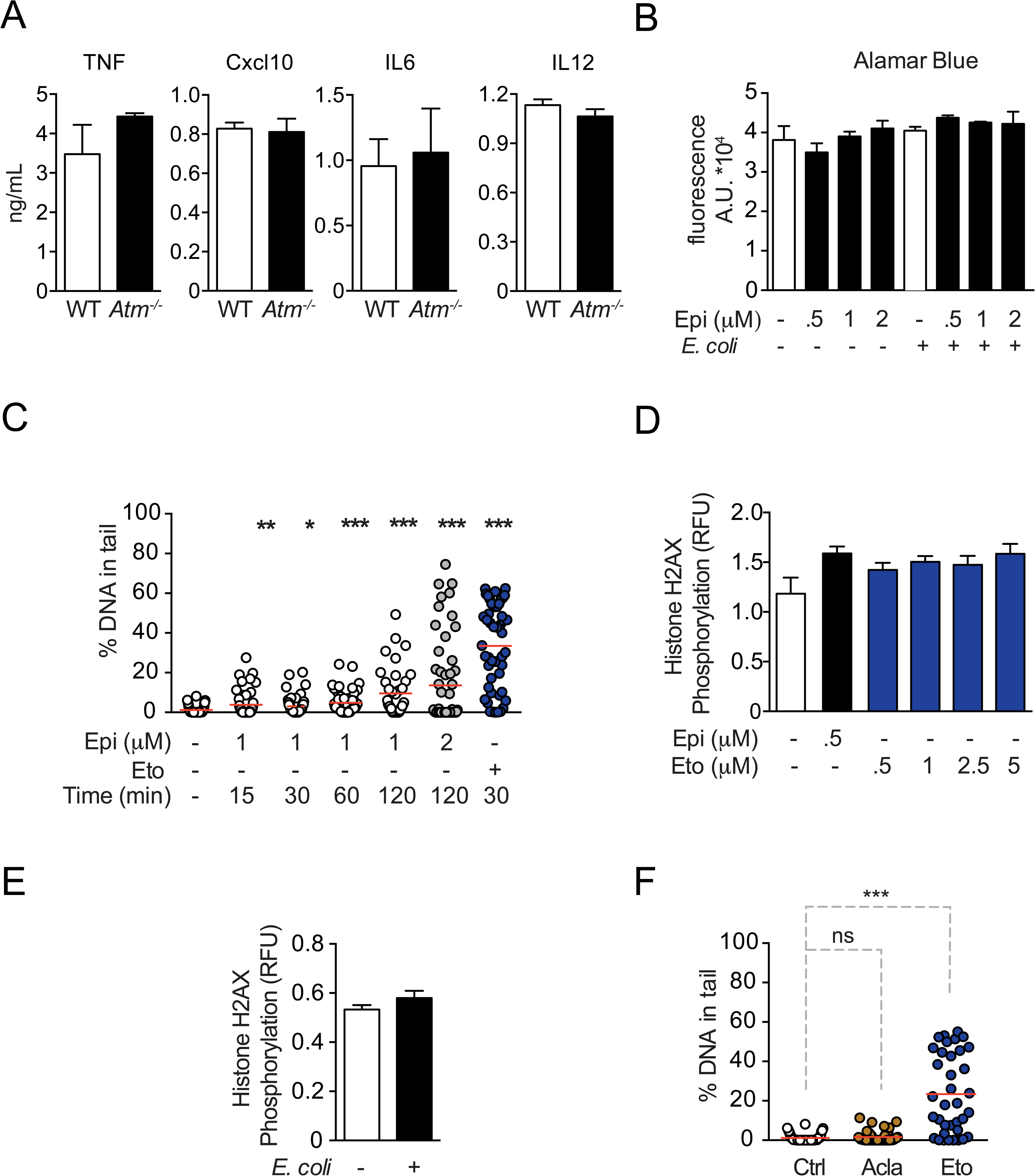
Control experiments for cytokine regulation by Epirubicin (Epi) and characterization of DNA damage. **A**, Following stimulation with *E. coli*, WT and *Atm*-/- macrophages secrete similar amounts of the cytokines TNF, IL12, IL6 and Cxcl10, as observed by ELISA; **B**, Cell number quantification using Alamar Blue fluorescence in unstimulated macrophages or macrophages stimulated with *E. coli*, both treated with various doses of Epi; **C**, Cytokine secretion of TNF, IL12, IL6 and Cxcl10, quantified by ELISA, is only detected in cells stimulated with *E. coli* and is not detected in unstimulated cells left untreated or treated with 2µM of Epi; **D**, Percentage of broken DNA in the comet tail as quantified by Comet Assay in THP-1 cells treated for 15, 30, 60 and 120 min with Epi or Etoposide (Eto); **E**, H2AX phosphorylation was quantified by ELISA in macrophages treated with various doses of Epi and Eto; **F**, Similar to **E** but the macrophages were either stimulated with *E. coli* for 4h our left unstimulated; **G**, Similar to **D** but the treatments were for 120 min with 1µM of Aclarubicin (Acla) and Eto. The assays show arithmetic means and standard deviations of technical replicates from one representative animal of at least three independent animals tested. p < 0.05 (*); p < 0.01 (**); p < 0.001 (***); p < 0.0001 (****).

**Supplementary Figure 2.**
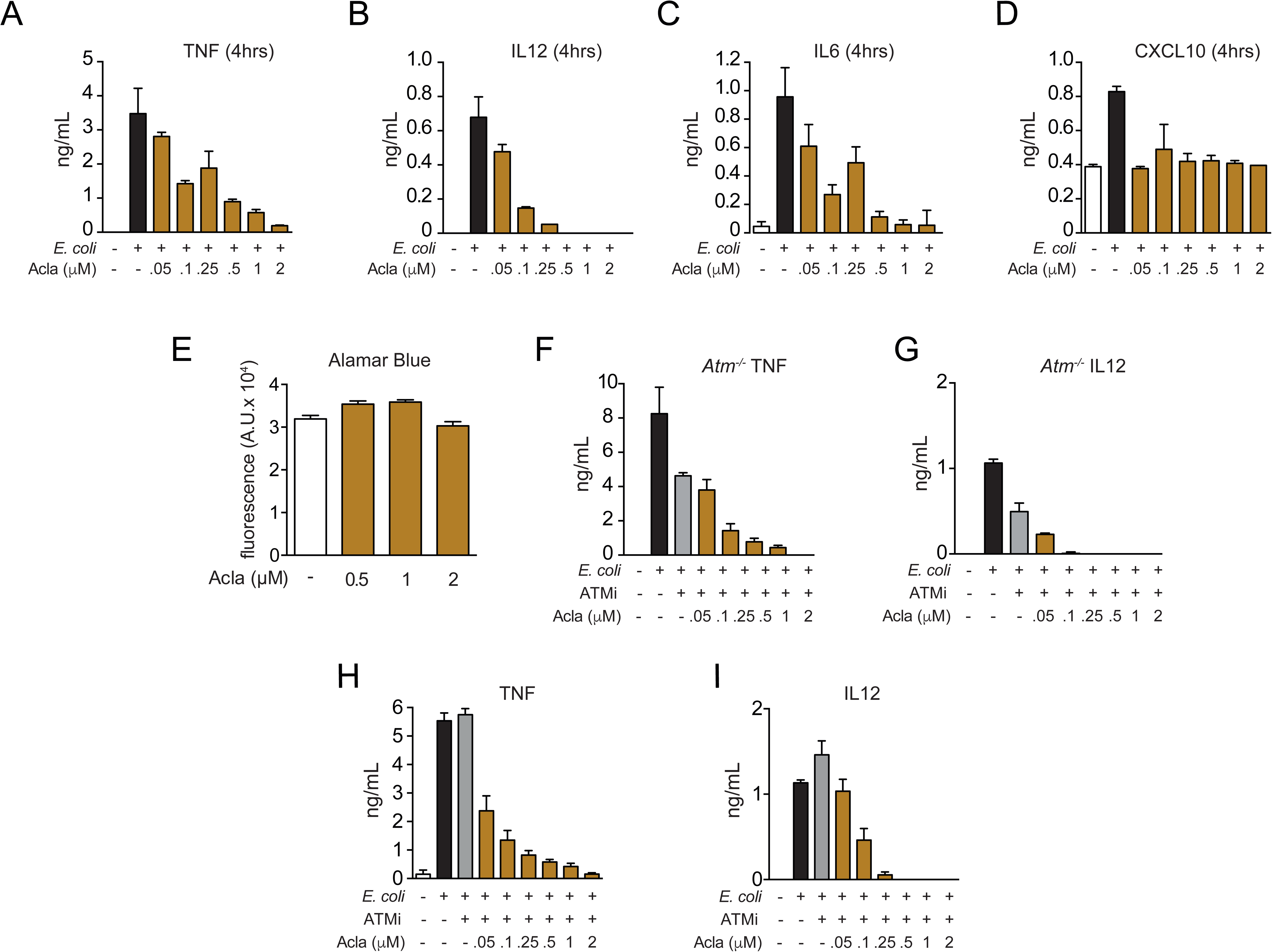
Similarly to Epirubicin (Epi), Aclarubicin (Acla) regulates cytokine secretion independently of ATM. **A** to **D**, Cytokine secretion of TNF, IL12, IL6 and Cxcl10 in macrophages was quantified by ELISA following *E. coli* challenge in the presence of various doses of Acla; **E**, Cell number quantification using Alamar Blue fluorescence in unstimulated macrophages treated with various doses of Acla; **F**, TNF and **G**, IL12 secretion was quantified by ELISA following *E. coli* challenge in the presence of various doses of Acla in *Atm*-/- macrophages; **H**, TNF and **I**, IL12 secretion was quantified by ELISA following *E. coli* challenge in macrophages treated with various doses of Acla and the ATM inhibitor KU-55933. The assays show arithmetic means and standard deviations of technical replicates from one representative animal of at least three independent animals tested. p < 0.05 (*); p < 0.01 (**); p < 0.001 (***); p < 0.0001 (****).

**Supplementary Figure 3.**
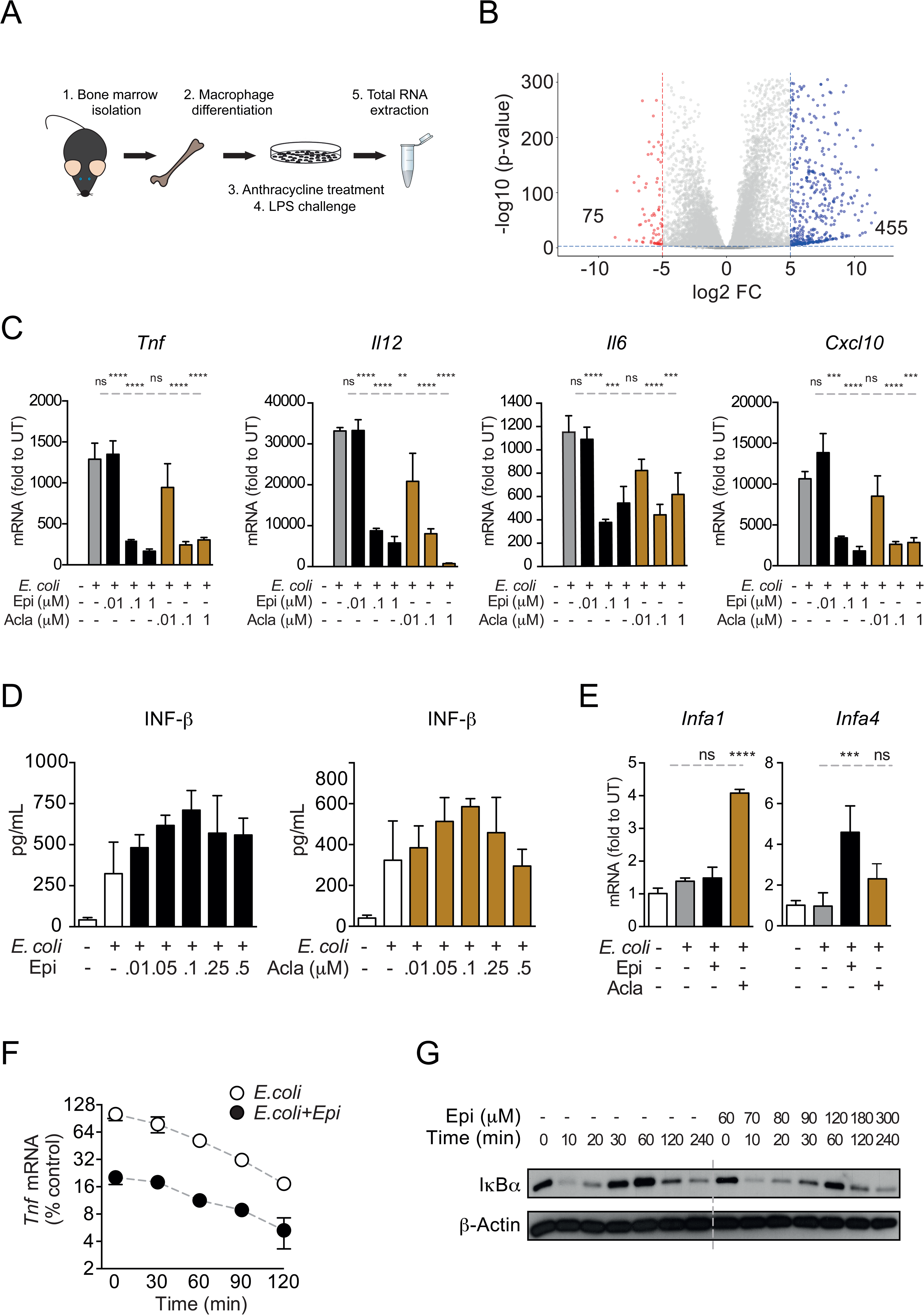
Testing specificity of transcriptional regulation by anthracyclines. **A**, RNAseq analysis workflow; **B**, Volcano plot of LPS-regulated genes from the RNAseq; **C**, Gene expression was quantified by quantitative RT-PCR in macrophages following *E. coli* challenge and treated with various doses of Epirubicin (Epi) and Aclarubicin (Acla); results were normalized to expression in untreated conditions (UT); **D**, Interferon beta secretion was quantified by ELISA following *E. coli* challenge in macrophages treated with various doses of Epi (left) and Acla (right); **E**, Gene expression was quantified as in C; **F**, Analysis of TNF mRNA stability in macrophages treated with 2µM of Epi for 1h or left untreated; Actinomycin D was added 2h after *E. coli* stimulation; **G**, I*κ*Bα degradation kinetics in macrophages following TNF challenge in the absence or presence of 2µM of Epi for 1h at the time of the challenge.

**Supplementary Table 1.** Functional analysis of Epirubicin (Epi) and Aclarubicin (Acla) repressed genes. DAVID functional analysis of the RNAseq data in macrophages stimulated with LPS for 4h; the table shows representative clusters of genes downregulated by Epi or Acla.

Cluster Enrichment Score: 7.128357433869625
| Category | Term |
| --- | --- |
| UniProt Keywords | Inflammatory response |
| UniProt Keywords | Immunity |
| UniProt Keywords | Innate immunity |
| GO Biological Process | Immune system process |
| GO Biological Process | innate immune response |

Cluster Enrichment Score: 6.570030482177336
| Category | Term |
| --- | --- |
| KEGG PATHWAY | Cytokine-cytokine receptor interaction |
| GO Biological Process | Immune response |
| UniProt Keywords | Cytokine |
| GO Biological Process | Cytokine activity |

Cluster Enrichment Score: 1.944662280964713
| Category | Term |
| --- | --- |
| INTERPRO | Tumour necrosis factor |
| SMART | TNF |
| INTERPRO | Tumour necrosis factor, conserved site |
| INTERPRO | Tumour necrosis factor alpha/beta/c |
| Go Molecular Function | Tumor necrosis factor receptor binding |
| INTERPRO | Tumour necrosis factor-like domain |

Cluster Enrichment Score: 26.374073343937372
| Category | Term |
| --- | --- |
| UniProt Keywords | Immunity |
| GO Biological Process | Immune system process |
| UniProt Keywords | Innate immunity |
| GO Biological Process | Innate immune response |

Cluster Enrichment Score: 9.853989037368624
| Category | Term |
| --- | --- |
| GO Biological Process | Immune response |
| KEGG PATHWAY | Cytokine-cytokine receptor interaction |
| UniProt Keywords | Cytokine |
| Go Molecular Function | Cytokine activity |

Cluster Enrichment Score: 3.8322404478340935
| Category | Term |
| --- | --- |
| GO Biological Process | Cellular response to interferon-gamma |
| GO Biological Process | Cell chemotaxis |
| GO Biological Process | Chemotaxis |
| UniProt Keywords | Chemotaxis |
| GO Biological Process | Chemokine-mediated signaling pathway |
| INTERPRO | CC chemokine, conserved site |
| INTERPRO | Chemokine interleukin-8-like domain |
| GO Biological Process | Neutrophil chemotaxis |
| Go Molecular Function | Chemokine activity |
| SMART | SCY |
| GO Biological Process | Lymphocyte chemotaxis |
| GO Biological Process | Monocyte chemotaxis |
| GO Biological Process | Positive regulation of natural killer cell chemotaxis |
| Go Molecular Function | CCR chemokine receptor binding |
| GO Biological Process | Macrophage chemotaxis |
| GO Biological Process | Eosinophil chemotaxis |

Epi
Acla

**Supplementary Figure 4.**
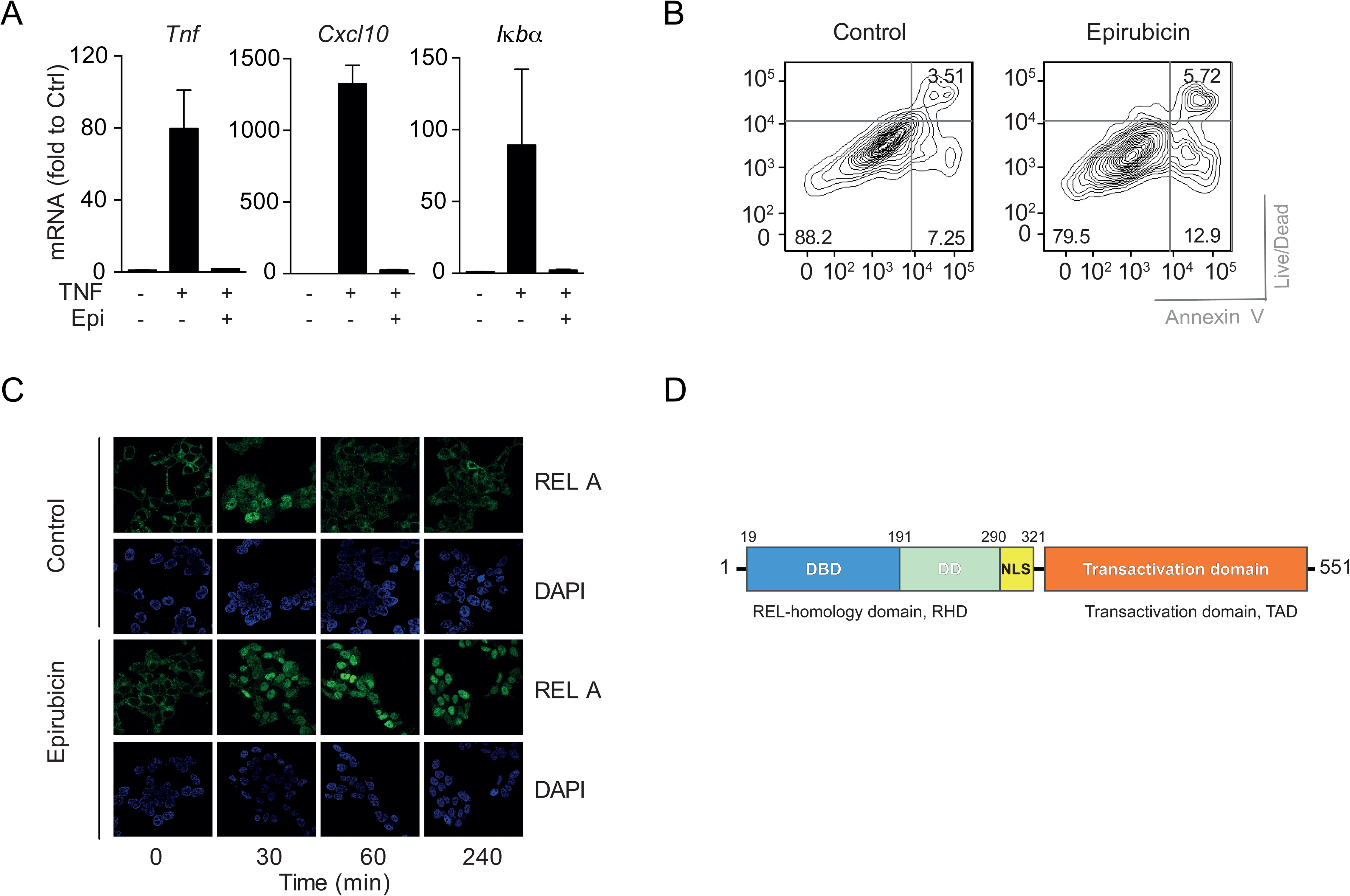
Epirubicin (Epi) modulates NF-*κ*B activity in HEK293 cells. **A**, TNF, Cxcl10 and IKBA mRNA levels in HEK293 cells challenged with TNF for 4 hours and treated with Epi; **B**, HEK293 cells were left untreated (control) or exposed to Epi for 16 hours. Cell viability was evaluated by flow cytometry based on Live/Dead and Annexin V staining; **C**, RelA immunolocalization in HEK293 cells left untreated (control) or exposed to 5μM of Epi for 1 hour, at the indicated time points, followed by TNF challenge. **D**, Schematic representation of RelA sequence with functional domains.

**Supplementary Figure 5.**
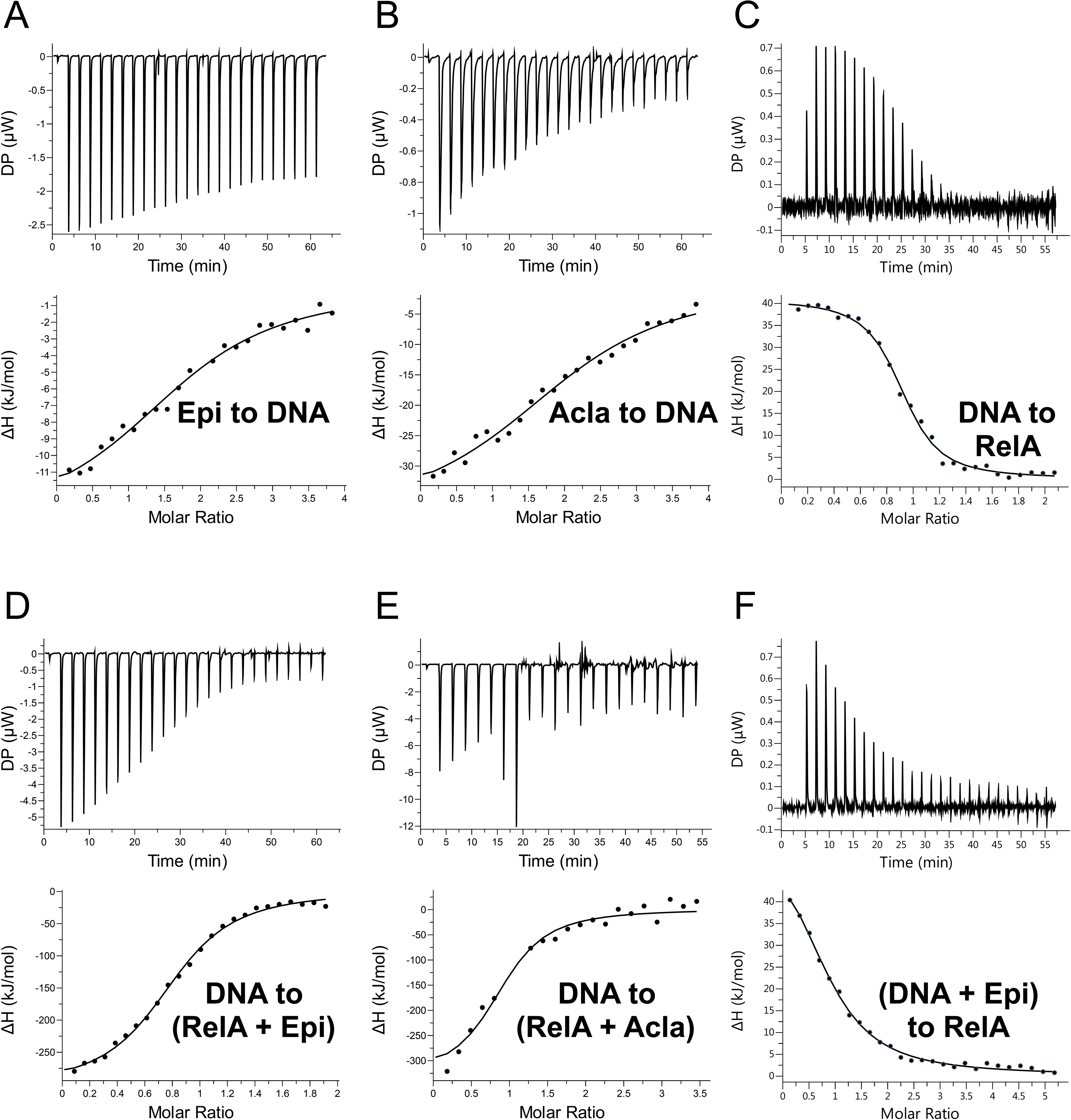
ITC binding measurements. **A**, 500μΜ Epirubicin (Epi) titrated to 25μΜ of duplex 14-mer *κ*B-33 DNA; **B**, 500μΜ Aclarubicin (Acla) titrated to 25μΜ of duplex 14-mer *κ*B-33 DNA; **C**, 125μΜ duplex 14-mer *κ*B-33 DNA titrated to 25μΜ of RelA; **D**, 125μΜ duplex 14-mer *κ*B-33 DNA titrated to 25μΜ RelA with 200μΜ Epi; **E**, 125μΜ duplex 14-mer *κ*B-33 DNA titrated to 25μΜ RelA with 200μΜ Acla and, **F**, 125μM duplex 14-mer *κ*B-33 DNA with 250μM Epirubicin titrated to 25μΜ of RelA.

**Supplementary Figure 6.**
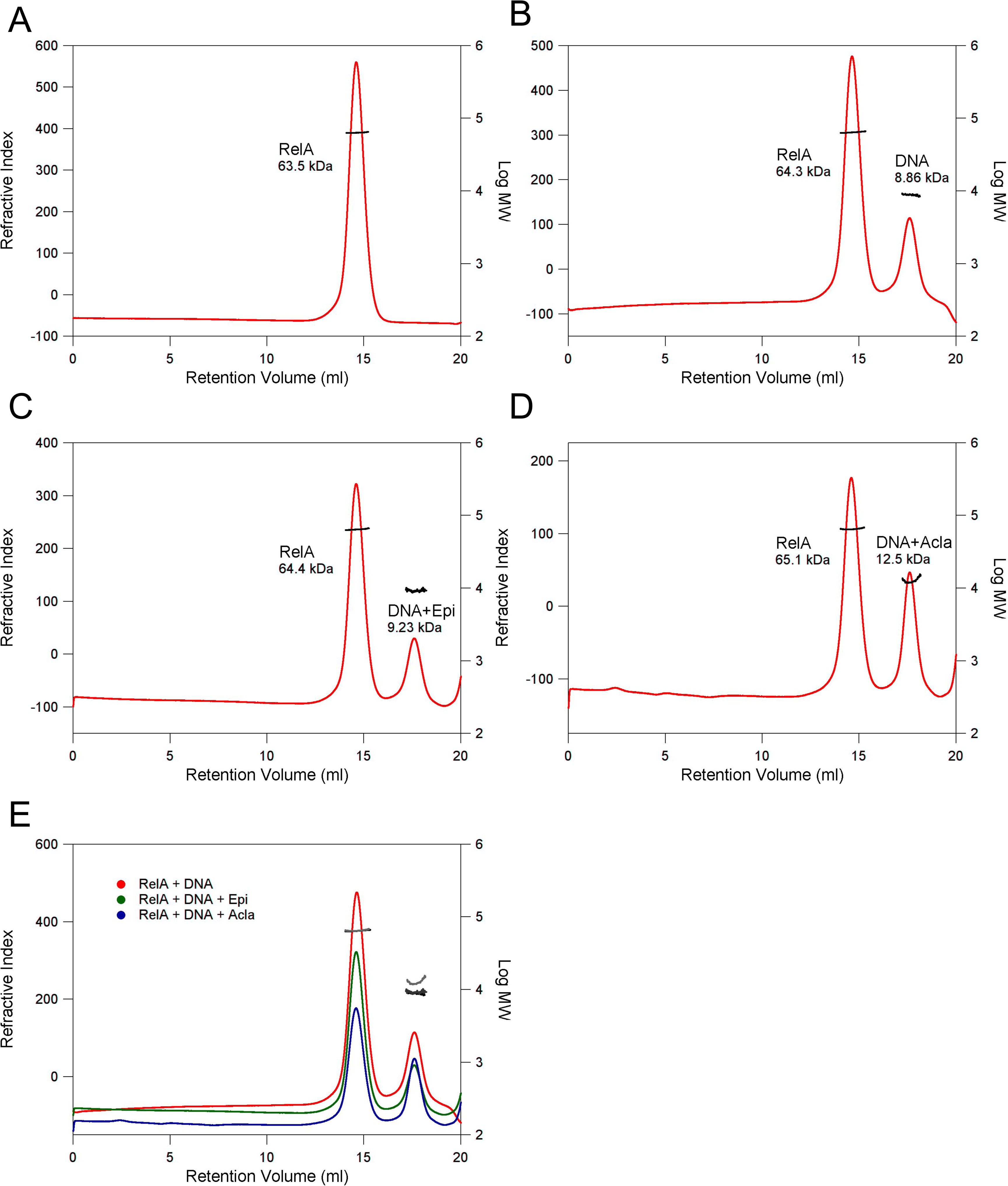
Determination of the molecular weight (MW) of complexes using size exclusion chromatography (SEC) in combination with static light scattering (SLS). **A,** RelA_19-291_; **B,** RelA_19-291_ with *κ*B-33 DNA (1:2 RelA:DNA); **C,** RelA_19-291_ with *κ*B-33 DNA and Epi (1:2:6 RelA:DNA:Epi); **D,** RelA_19-291_ with *κ*B-33 DNA and Acla (1:2:6 RelA:DNA:Acla); The refractive index (red) and right-angle light scattering (not shown) signals were monitored and used to determine the molecular weights (black). **E,** Superimposition of **B-D.** The refractive index and MW are shown in red and black (B); green and dark grey (C); blue and light grey (D), respectively.

**Supplementary Table 2.** Molecular weight values of RelA_19-291_, *κ*B-33 DNA, *κ*B-33 DNA + Epirubicin (Epi) and *κ*B-33 DNA + Aclarubicin (Acla) using SEC in combination with SLS (Supplementary Figure 6).

Epi Acla
| **Experiment** | **MW<sub>exp</sub>**<br>**RelA (kDa)** | **MW<sub>calc</sub>**<br>**RelA (kDa)** | **MW<sub>exp</sub>**<br>**DNA (kDa)** | **MW<sub>calc</sub>**<br>**DNA (kDa)** | **MW<sub>calc</sub>**<br>**Compounds**<br>**(kDa)** |
| --- | --- | --- | --- | --- | --- |
| **RelA** | 63.5 | 62.3 |  |  |  |
| **DNA** |  |  | 8.86 | 8.53 |  |
| **Epi** |  |  |  |  | 0.54 |
| **Acla** |  |  |  |  | 0.81 |
| **RelA + DNA (1:2)** | 64.3 |  | 8.86 |  |  |
| **RelA + DNA + Epi (1:2:6)** | 64.4 |  | 9.23 |  |  |
| **RelA + DNA + Acla (1:2:6)** | 65.1 |  | 12.5 |  |  |

**Supplementary Figure 7.**
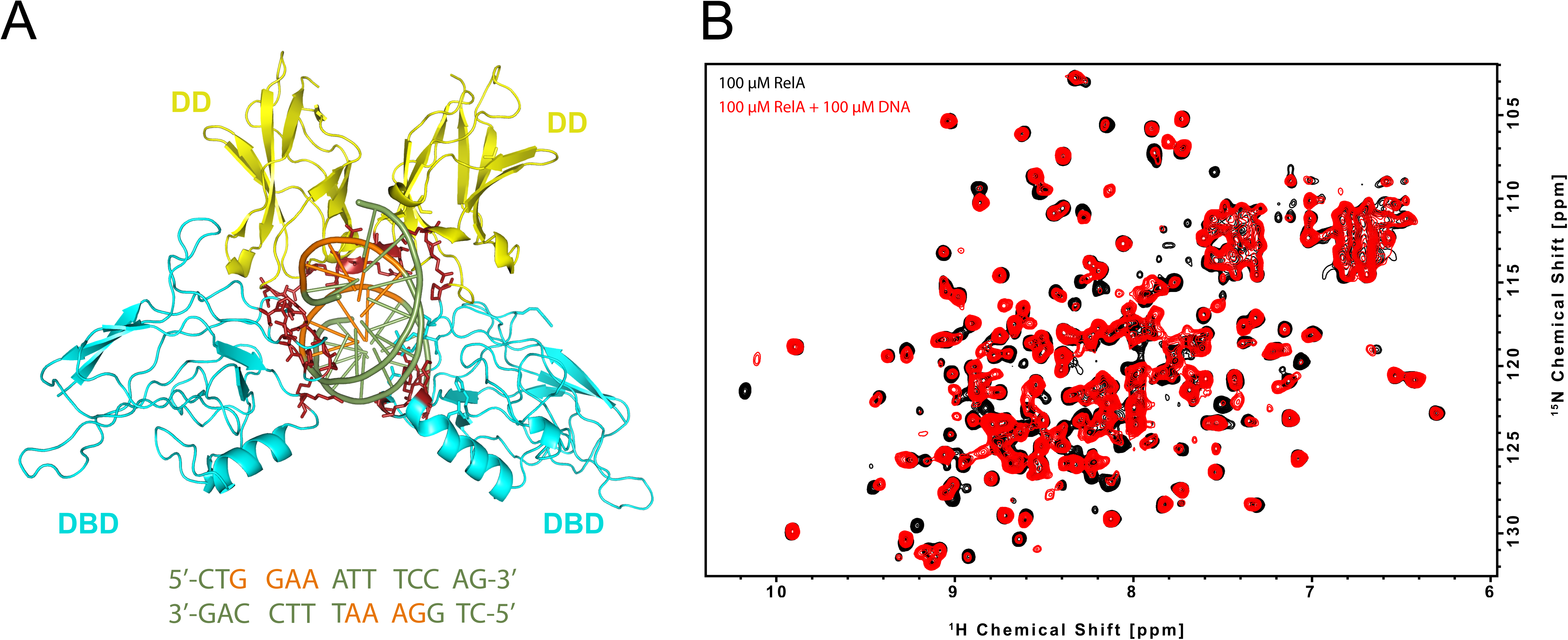
RelA RHD binding to *κ*B-33 promoter DNA **A**, 3D structure of RelA RHD in complex with *κ*B-33 promoter DNA (PDB 1RAM). The DD domain and DBD are shown in yellow and cyan, respectively. The residues that interact with the DNA are represented in dark red. The DNA is shown in green with nucleotides contacting the protein colored orange. The sequence of the 14-mer *κ*B-33 DNA derived from the 18-mer used for this crystal structure is shown below; **B**, NMR binding studies on RelA RHD domain with the 14-mer *κ*B-33 DNA. 2D ^1^H,^15^N correlation spectra of 100μΜ ^2^H, ^15^N RelA_19-291_ dimer recorded without (black) and with 100μΜ DNA duplex (red) (800MHz, 20 °C, 16 scans).

**Supplementary Figure 8.**
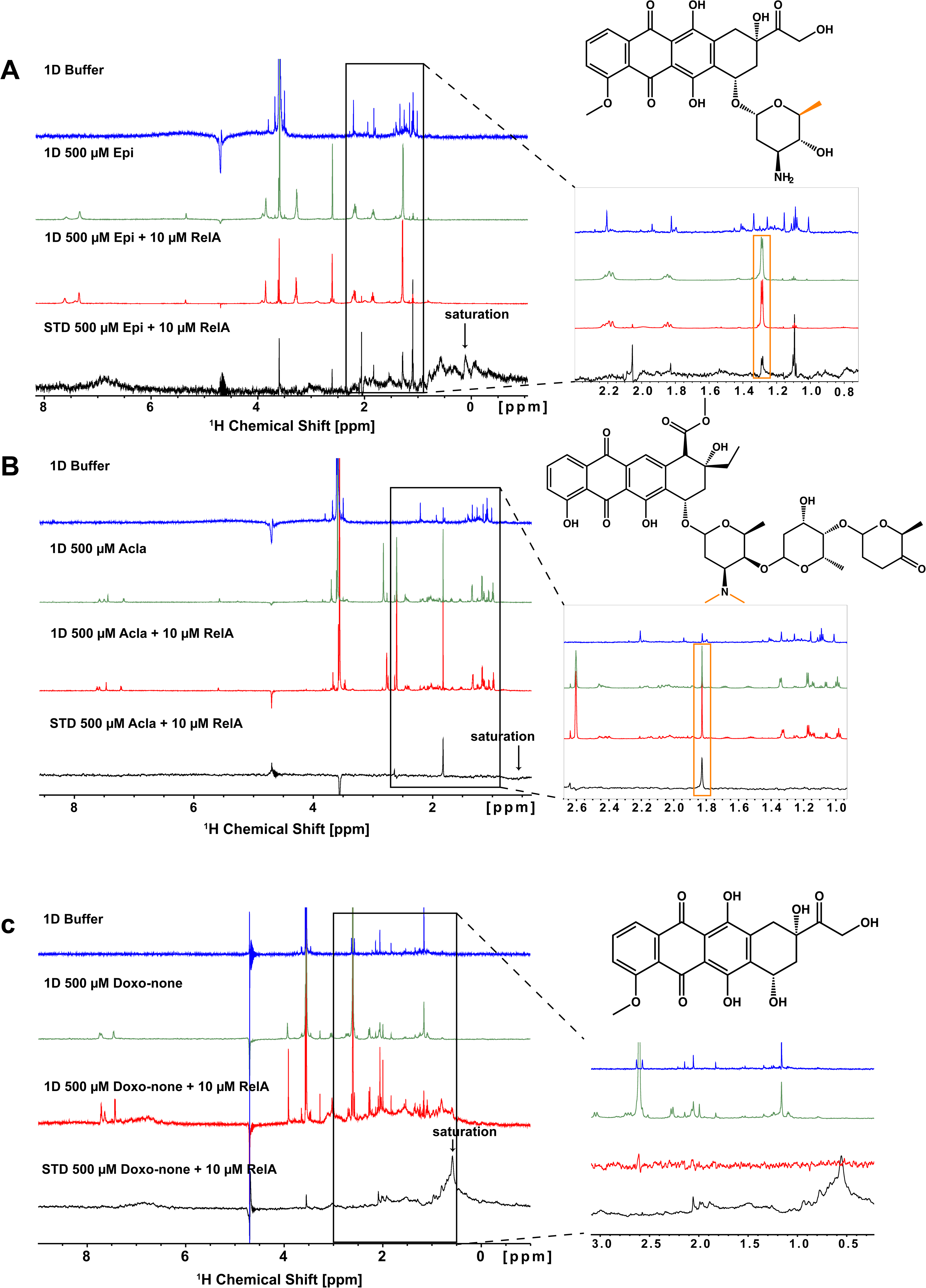
STD-NMR experiments to test the interaction of RelA with the anthracyclines Epirubicin (Epi), Aclarubicin (Acla) and Doxorubicinone (Doxo-none). Epi and Acla interact through the methyl groups(s) in the sugar rings with RelA. Doxo-none lacks a sugar ring and does not interact with RelA. **A,** Epi, **B,** Acla and **C,** Doxo-none. The blue spectra correspond to the reference buffer. The green spectra are recorded with 500μM of each compound in 100mM d11-Tris-HCl pH 7.5, 75mM NaCl and 10% D_2_O buffer (600MHz, 20°C, 128 scans). Red spectra correspond to the 1D spectra of the compounds with protein in buffer. The black spectra correspond to the STD spectra of the 10μM RelA with the 500μM anthracyclines. On the right side of each set of spectra, a zoomed view is shown highlighting the methyl groups showing STD signals (see above compound structure). In all STD experiments, the arrow indicates the irradiation region (0.05ppm) (600MHz, 20°C, 800 scans).

**Supplementary Figure 9.**
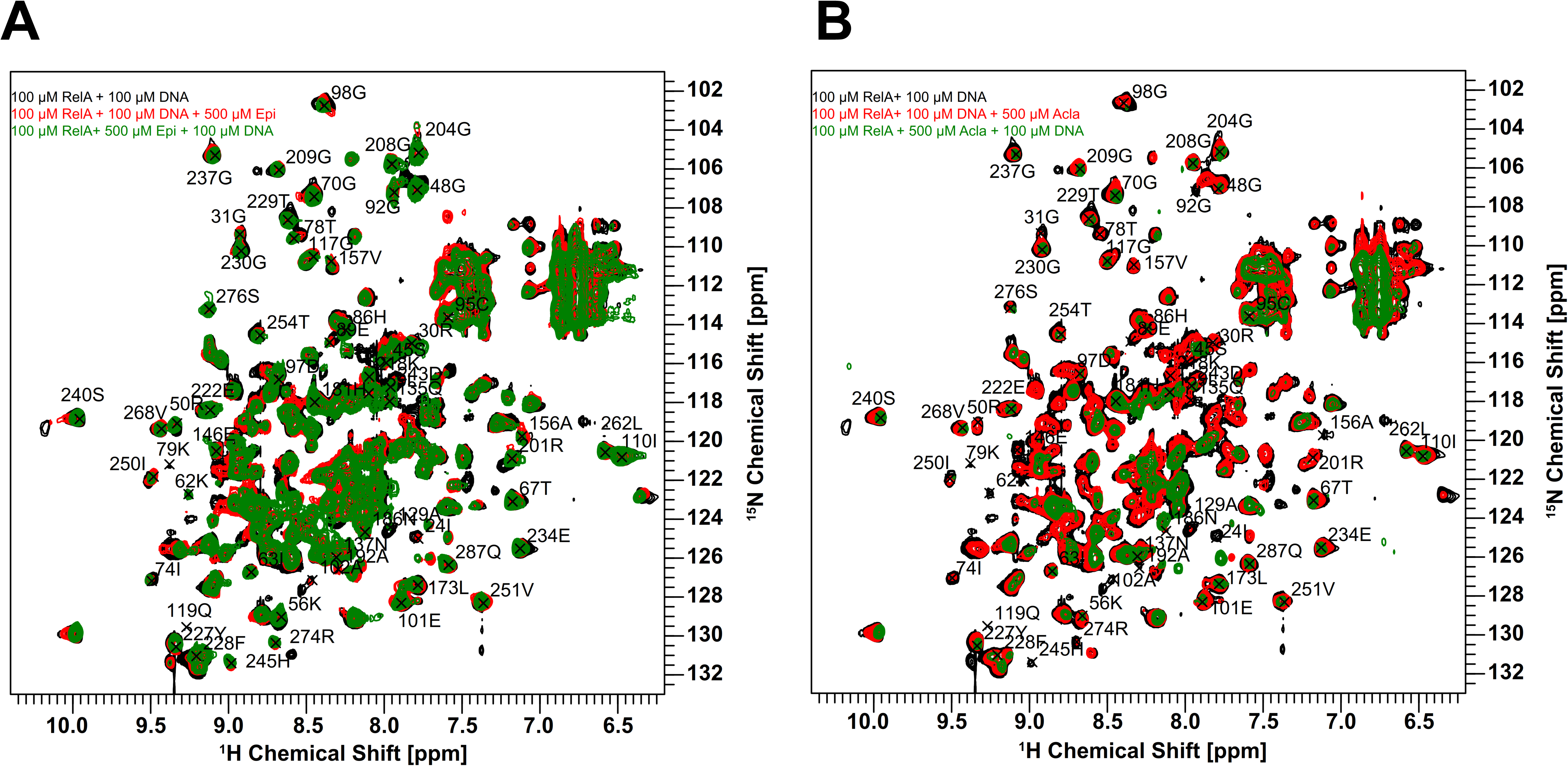
Comparison of the effect of the order of addition in NMR binding studies of Epirubicin (Epi) and Aclarubicin (Acla) with RelA and *κ*B-33 DNA. Superposition of 2D ^1^H,^15^N correlation spectra of 100μΜ ^2^H, ^15^N RelA_19-291_ with 100μΜ DNA (black) with **A,** ^2^H, ^15^N 100μΜ RelA_19-291_ with 100μΜ DNA and 100μΜ Epi (red) and ^2^H, ^15^N 100μΜ RelA_19-291_ with 500μΜ Epi and 100 μΜ DNA (green); and with **B,** ^2^H, ^15^N 100μΜ RelA_19-291_ with 100μΜ DNA and 500μΜ Acla (red) and ^2^H, ^15^N 100μΜ RelA_19-291_ with 500μΜ Acla and 100μΜ DNA (green) (800MHz, 20°C, 16 scans). The addition order of anthracyclines and DNA to RelA leads to spectra that are not completely superimposable suggesting that the complexes are different. Note that Aclarubicin causes protein precipitation, leading to larger spectral differences, especially when it is added directly to the protein.

**Supplementary Figure 10.**
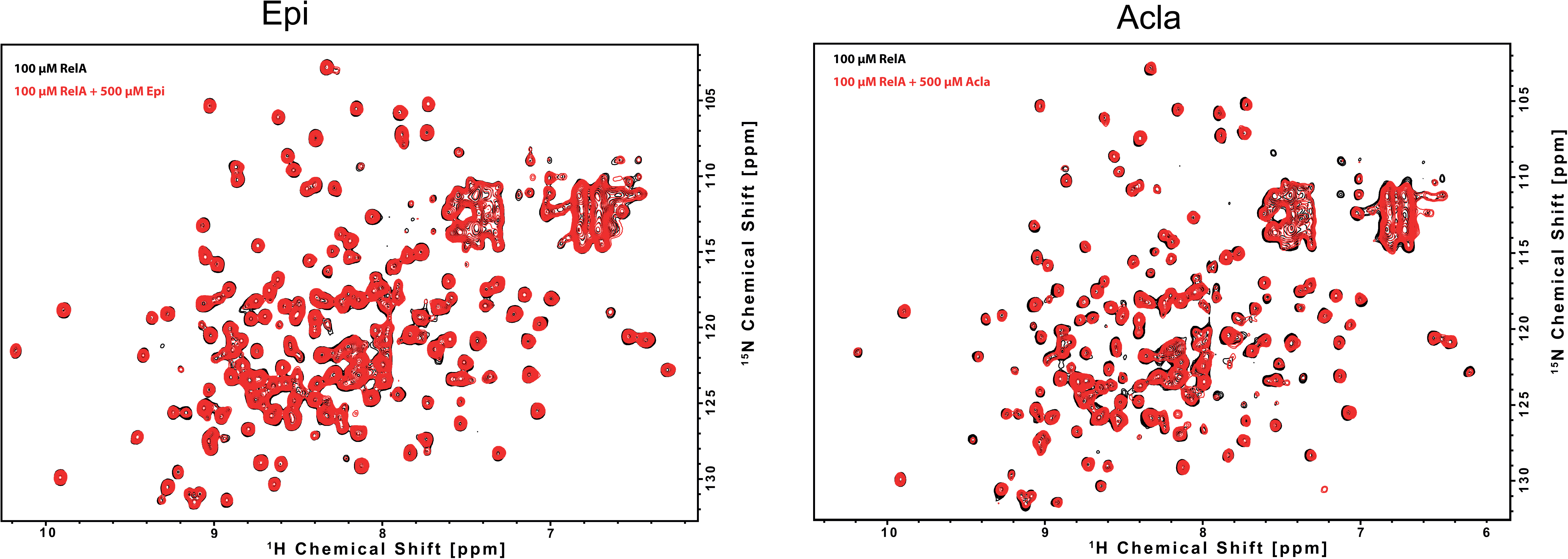
NMR binding studies on RelA_19-291_ with Epirubicin (Epi) and Aclarubicin (Acla). Superposition of 2D ^1^H,^15^N NMR correlation spectra of 100 μΜ ^2^H, ^15^N RelA_19-291_ recorded without (black) and with (left) 500μΜ of Epi (red) or (right) 500μΜ of Acla (red) (800 MHz, 20°C, 16 scans).

## References

De Almeida, S.F., Grosso, A.R., Koch, F., Fenouil, R., Carvalho, S., Andrade, J., Levezinho, H., Gut, M., Eick, D., Gut, I., et al. (2011). Splicing enhances recruitment of methyltransferase HYPB/Setd2 and methylation of histone H3 Lys36. Nat. Struct. Mol. Biol.

Anrather, J., Csizmadia, V., Soares, M.P., and Winkler, H. (1999). Regulation of NF-κB RelA phosphorylation and transcriptional activity by p21(ras) and protein kinase Cζ in primary endothelial cells. J. Biol. Chem.

Van Antwerp, D.J., Martin, S.J., Kafri, T., Green, D.R., and Verma, I.M. (1996). Suppression of TNF-α-induced apoptosis by NF-κB. Science (80-.).

Arlt, A., Vorndamm, J., Breitenbroich, M., Fölsch, U.R., Kalthoff, H., Schmidt, W.E., and Schäfer, H. (2001). Inhibition of NF-κB sensitizes human pancreatic carcinoma cells to apoptosis induced by etoposide (VP16) or doxorubicin. Oncogene.

Baldwin, A.S. (2001). Series Introduction: The transcription factor NF-κB and human disease. J. Clin. Invest.

Banáth, J.P., and Olive, P.L. (2003). Expression of phosphorylated histone H2AX as a surrogate of cell killing by drugs that create DNA double-strand breaks. Cancer Res.

Beg, A.A., and Baldwin, A.S. (1993). The IκB proteins: Multifunctional regulators of Rel/NF-κB transcription factors. Genes Dev.

Bergqvist, S., Alverdi, V., Mengel, B., Hoffmann, A., Ghosh, G., and Komives, E.A. (2009). Kinetic enhancement of NF-κB·DNA dissociation by IκBα. Proc. Natl. Acad. Sci. U. S. A.

Borghesani, P.R., Alt, F.W., Bottaro, A., Davidson, L., Aksoy, S., Rathbun, G.A., Roberts, T.M., Swat, W., Segal, R.A., and Gu, Y. (2000). Abnormal development of Purkinje cells and lymphocytes in Atm mutant mice. Proc. Natl. Acad. Sci. U. S. A.

Campbell, K.J., Rocha, S., and Perkins, N.D. (2004). Active repression of antiapoptotic gene expression by RelA(p65) NF-κB. Mol. Cell.

Caporossi, D., Porfirio, B., Nicoletti, B., Palitti, F., Degrassi, F., De Salvia, R., and Tanzarella, C. (1993). Hypersensitivity of lymphoblastoid lines derived from ataxia telangiectasia patients to the induction of chromosomal aberrations by etoposide (VP-16). Mutat. Res. - Fundam. Mol. Mech. Mutagen.

Chaires, J.B. (2015). A small molecule - DNA binding landscape. Biopolymers.

Chang, J.T., and Nevins, J.R. (2006). GATHER: A systems approach to interpreting genomic signatures. Bioinformatics.

Chen, L.F., and Greene, W.C. (2004). Shaping the nuclear action of NF-κB. Nat. Rev. Mol. Cell Biol.

Chen, Y.Q., Ghosh, S., and Ghosh, G. (1998). A novel DNA recognition mode by the NF-κb p65 homodimer. Nat. Struct. Biol.

Chen, Y.Q., Sengchanthalangsy, L.L., Hackett, A., and Ghosh, G. (2000). NF-κB p65 (RelA) homodimer uses distinct mechanisms to recognize DNA targets. Structure.

Colaço, H.G., Barros, A., Neves-Costa, A., Seixas, E., Pedroso, D., Velho, T., Willmann, K.L., Faisca, P., Grabmann, G., Yi, H.S., et al. (2021). Tetracycline Antibiotics Induce Host-Dependent Disease Tolerance to Infection. Immunity.

Delaglio, F., Grzesiek, S., Vuister, G.W., Zhu, G., Pfeifer, J., and Bax, A. (1995). NMRPipe: A multidimensional spectral processing system based on UNIX pipes. J. Biomol. NMR.

Dobin, A., Davis, C.A., Schlesinger, F., Drenkow, J., Zaleski, C., Jha, S., Batut, P., Chaisson, M., and Gingeras, T.R. (2013). STAR: Ultrafast universal RNA-seq aligner. Bioinformatics.

Eom, Y.W., Kim, M.A., Park, S.S., Goo, M.J., Kwon, H.J., Sohn, S., Kim, W.H., Yoon, G., and Choi, K.S. (2005). Two distinct modes of cell death induced by doxorubicin: Apoptosis and cell death through mitotic catastrophe accompanied by senescence-like phenotype. Oncogene.

Ewels, P., Magnusson, M., Lundin, S., and Käller, M. (2016). MultiQC: Summarize analysis results for multiple tools and samples in a single report. Bioinformatics.

Figueiredo, N., Chora, A., Raquel, H., Pejanovic, N., Pereira, P., Hartleben, B., Neves-Costa, A., Moita, C., Pedroso, D., Pinto, A., et al. (2013). Anthracyclines induce DNA damage response-mediated protection against severe sepsis. Immunity.

Frederick, C.A., Williams, L.D., Ughetto, G., van der Marel, G.A., van Boom, H.J., Rich, A., and Wang, A.H.J. (1990). Structural Comparison of Anticancer Drug-DNA Complexes: Adriamycin and Daunomycin. Biochemistry.

Furusawa, H., Nakayama, H., Funasaki, M., and Okahata, Y. (2016). Kinetic characterization of small DNA-binding molecules interacting with a DNA strand on a quartz crystal microbalance. Anal. Biochem.

Ghosh, S., and Baltimore, D. (1990). Activation in vitro of NF-κB” by phosphorylation of its inhibitor IκB”. Nature.

Hande, K.R. (1998). Clinical applications of anticancer drugs targeted to topoisomerase II. Biochim. Biophys. Acta - Gene Struct. Expr.

Hayden, M.S., and Ghosh, S. (2008). Shared Principles in NF-κB Signaling. Cell.

Ho, W.C., Dickson, K.M., and Barker, P.A. (2005). Nuclear factor-κB induced by doxorubicin is deficient in phosphorylation and acetylation and represses nuclear factor-κB-dependent transcription in cancer cells. Cancer Res.

Hochrainer, K., Racchumi, G., and Anrather, J. (2007). Hypo-phosphorylation leads to nuclear retention of NF-κB p65 due to impaired IκBα gene synthesis. FEBS Lett.

Huang, B., Yang, X.D., Lamb, A., and Chen, L.F. (2010). Posttranslational modifications of NF-κB: Another layer of regulation for NF-κB signaling pathway. Cell. Signal.

Janssens, S., Tinel, A., Lippens, S., and Tschopp, J. (2005). PIDD Mediates NF-κB activation in response to DNA damage. Cell.

Katenkamp, U., Stutter, E., Petri, I., Gollmick, F.A., and Berg, H. (1983). Interaction of anthracycline antibiotics with biopolymers VIII. binding parameters of aclacinomycin A to Dna. J. Antibiot. (Tokyo).

Kawai, T., and Akira, S. (2007). Signaling to NF-κB by Toll-like receptors. Trends Mol. Med.

Köse-Vogel, N., Stengel, S., Gardey, E., Kirchberger-Tolstik, T., Reuken, P.A., Stallmach, A., and Bruns, T. (2020). Transcriptional Suppression of the NLRP3 Inflammasome and Cytokine Release in Primary Macrophages by Low-Dose Anthracyclines. Cells 9.

Li, Y., Carty, M.P., Oakley, G.G., Seidman, M.M., Medvedovic, M., and Dixon, K. (2001). Expression of ATM in ataxia telangiectasia fibroblasts rescues defects in DNA double-strand break repair in nuclear extracts. Environ. Mol. Mutagen.

Liu, J., Zheng, H., Tang, M., Ryu, Y.C., and Wang, X. (2008). A therapeutic dose of doxorubicin activates ubiquitin-proteasome system-mediated proteolysis by acting on both the ubiquitination apparatus and proteasome. Am. J. Physiol. - Hear. Circ. Physiol.

Love, M.I., Huber, W., and Anders, S. (2014). Moderated estimation of fold change and dispersion for RNA-seq data with DESeq2. Genome Biol.

Maréchal, A., and Zou, L. (2013). DNA damage sensing by the ATM and ATR kinases. Cold Spring Harb. Perspect. Biol.

Medzhitov, R., and Horng, T. (2009). Transcriptional control of the inflammatory response. Nat. Rev. Immunol.

Millour, J., De Olano, N., Horimoto, Y., Monteiro, L.J., Langer, J.K., Aligue, R., Hajji, N., and Lam, E.W.F. (2011). ATM and p53 regulate FOXM1 expression via E2F in breast cancer epirubicin treatment and resistance. Mol. Cancer Ther.

Moura-alves, P., Neves-costa, A., Raquel, H., Pacheco, T.R., Almeida, B.D., Oliveira, M., Gama-carvalho, M., Rodrigues, R., Cadima-couto, I., Hacohen, N., et al. (2011). An shRNA-Based Screen of Splicing Regulators Identifies SFRS3 as a Negative Regulator of IL-1b Secretion. 6, 1–10.

Mukherjee, S.P., Borin, B., Quintas, P.O., and Dyson, H.J. (2016). NMR characterization of a 72 kDa transcription factor using differential isotopic labeling. Protein Sci.

Mulero, M.C., Huang, D. Bin, Thien Nguyen, H., Wang, V.Y.F., Li, Y., Biswas, T., and Ghosh, G. (2017). DNA-binding affinity and transcriptional activity of the RelA homodimer of nuclear factor κb are not correlated. J. Biol. Chem.

Neves-Costa, A., and Moita, L.F. (2017). Modulation of inflammation and disease tolerance by DNA damage response pathways. FEBS J. 284.

Nitiss, J.L. (2009). Targeting DNA topoisomerase II in cancer chemotherapy. Nat. Rev. Cancer.

Pang, B., Qiao, X., Janssen, L., Velds, A., Groothuis, T., Kerkhoven, R., Nieuwland, M., Ovaa, H., Rottenberg, S., Van Tellingen, O., et al. (2013). Drug-induced histone eviction from open chromatin contributes to the chemotherapeutic effects of doxorubicin. Nat. Commun.

Pang, B., De Jong, J., Qiao, X., Wessels, L.F.A., and Neefjes, J. (2015). Chemical profiling of the genome with anti-cancer drugs defines target specificities. Nat. Chem. Biol.

Piret, B., Schoonbroodt, S., and Piette, J. (1999). The ATM protein is required for sustained activation of NF-κB following DNA damage. Oncogene.

Qiao, X., Van Der Zanden, S.Y., Wander, D.P.A., Borràs, D.M., Song, J.Y., Li, X., Duikeren, S. Van, Gils, N. Van, Rutten, A., Herwaarden, T. Van, et al. (2020). Uncoupling DNA damage from chromatin damage to detoxify doxorubicin. Proc. Natl. Acad. Sci. U. S. A.

Sattler, M., Schleucher, J., and Griesinger, C. (1999). Heteronuclear multidimensional NMR experiments for the structure determination of proteins in solution employing pulsed field gradients. Prog. Nucl. Magn. Reson. Spectrosc.

Skovsgaard, T. (1987). Pharmacodynamic aspects of aclarubicin with special reference to daunorubicin and doxorubicin. Eur. J. Haematol.

Stephens, M. (2017). False discovery rates: A new deal. Biostatistics.

Sun, S.C., Ganchi, P.A., Ballard, D.W., and Greene, W.C. (1993). NF-κB controls expression of inhibitor IκBα: Evidence for an inducible autoregulatory pathway. Science (80-.).

Tanaka, T., Yamaguchis, J., Shojis, K., and Nangakus, M. (2012). Anthracycline inhibits recruitment of hypoxia-inducible transcription factors and suppresses tumor cell migration and cardiac angiogenic response in the host. J. Biol. Chem.

Temperini, C., Messori, L., Orioli, P., Di Bugno, C., Animati, F., and Ughetto, G. (2003). The crystal structure of the complex between a disaccharide anthracycline and the DNA hexamer d(CGATCG) reveals two different binding sites involving two DNA duplexes. Nucleic Acids Res.

Tewey, K.M., Rowe, T.C., Yang, L., Halligan, B.D., and Liu, L.F. (1984). Adriamycin-induced DNA damage mediated by mammalian DNA topoisomerase II. Science (80-.).

Toledano, M.B., Ghosh, D., Trinh, F., and Leonard, W.J. (1993). N-terminal DNA- binding domains contribute to differential DNA-binding specificities of NF-kappa B p50 and p65. Mol. Cell. Biol.

Utsuno, K., and Tsuboi, M. (1997). Degree of DNA unwinding caused by the binding of aclacinomycin A. Chem. Pharm. Bull.

Vranken, W.F., Boucher, W., Stevens, T.J., Fogh, R.H., Pajon, A., Llinas, M., Ulrich, E.L., Markley, J.L., Ionides, J., and Laue, E.D. (2005). The CCPN data model for NMR spectroscopy: Development of a software pipeline. Proteins Struct. Funct. Genet.

Wander, D.P.A., van der Zanden, S.Y., van der Marel, G.A., Overkleeft, H.S., Neefjes, J., and Codeé, J.D.C. (2020). Doxorubicin and aclarubicin: Shuffling anthracycline glycans for improved anticancer agents. J. Med. Chem.

Wang, C.Y., Mayo, M.W., and Baldwin, A.S. (1996). TNF- and cancer therapy-induced apoptosis: Potentiation by inhibition of NF-κB. Science (80-.).

Wu, M., Lee, H., Bellas, R.E., Schauer, S.L., Arsura, M., Katz, D., Fitzgerald, M.J., Rothstein, T.L., Sherr, D.H., and Sonenshein, G.E. (1996). Inhibition of NF-κB/Rel induces apoptosis of murine B cells. EMBO J.

Wu, Z.H., Shi, Y., Tibbetts, R.S., and Miyamoto, S. (2006). Molecular linkage between the kinase ATM and NF-κB signaling in response to genotoxic stimuli. Science (80-.).

Yang, F., Kemp, C.J., and Henikoff, S. (2013). Doxorubicin enhances nucleosome turnover around promoters. Curr. Biol.

Zhang, O., Kay, L.E., Olivier, J.P., and Forman-Kay, J.D. (1994). Backbone 1H and 15N resonance assignments of the N-terminal SH3 domain of drk in folded and unfolded states using enhanced-sensitivity pulsed field gradient NMR techniques. J. Biomol. NMR.

Zunino, F., Pratesi, G., and Perego, P. (2001). Role of the sugar moiety in the pharmacological activity of anthracyclines: Development of a novel series of disaccharide analogs. Biochem. Pharmacol.

